# Paralytic, the *Drosophila* voltage-gated sodium channel, regulates proliferation of neural progenitors

**DOI:** 10.1101/803288

**Authors:** Beverly J. Piggott, Christian J. Peters, Ye He, Xi Huang, Susan Younger, Lily Yeh Jan, Yuh Nung Jan

## Abstract

Proliferating cells, typically considered “non-excitable,” nevertheless exhibit regulation by bioelectrical signals. Notably, voltage-gated sodium channels (VGSC) that are crucial for neuronal excitability, are also found in progenitors and upregulated in cancer. Here, we identify a role for VGSC in proliferation of *Drosophila* neuroblast (NB) lineages within the central nervous system. Loss of *paralytic (para)*, the sole gene that encodes *Drosophila* VGSC, reduces neuroblast progeny cell number. The type II neuroblast lineages, featuring a transit-amplifying intermediate neural progenitors (INP) population similar to that found in the developing human cortex, are particularly sensitive to *para* manipulation. Following a series of asymmetric divisions, INPs normally exit the cell cycle through a final symmetric division. Our data suggests that loss of *para* induces apoptosis in this population, whereas overexpression leads to an increase in INPs and overall neuroblast progeny cell numbers. These effects are cell autonomous and depend on Para channel activity. Reduction of Para not only affects normal NB development, but also strongly suppresses brain tumor mass, implicating a role for Para in cancer progression. To our knowledge, our studies are the first to identify a role for VGSC in neural progenitor proliferation. Elucidating the contribution of VGSC in proliferation will advance our understanding of bioelectric signaling within development and disease states.

## Introduction

Neuronal excitability is a defining feature of nervous systems. Voltage-gated ion channels mediate changes in electrical potential (voltage) across cell membranes, triggering second messenger and gene regulatory cascades. Generation of these bioelectric signals is crucial for neuronal excitability, but their involvement in neural development remains an open question. The formation of a properly functioning brain is a complex, temporal process where stem cells must balance self-renewal with differentiation to ensure the generation of enough neurons, with the correct identities, distribution and connectivity. Expression of voltage-gated ion channels is typically considered a post mitotic event, occurring within differentiated, excitable tissues. However, bioelectric signals govern biology in every living cell type where the asymmetric distribution of ions across the plasma membrane establishes a membrane potential (Vm). Vm is determined by the ion permeability and abundance of various ion channels, pumps, and exchangers within a given cell type. Throughout development, tissues typically thought to be nonexcitable are subjected to changes in Vm, governing wideranging cell behaviors including proliferation, migration, differentiation and death (McLaughlin and Levin 2018). The importance of bioelectric signals in non-excitable tissues is evident as channelopathies include diseases that affect embryonic patterning and development, with consequences as severe as limb and craniofacial abnormalities (McLaughlin and Levin 2018). Despite emerging evidence for bioelectric processes influencing brain development, the molecular and cellular basis for this is largely unknown.

The *Drosophila melanogaster* larval nervous system is a well-established model for elucidating mechanisms of neurogenesis (Doe 2008; Homem and Knoblich 2012; Homem et al. 2015; Farnsworth and Doe 2017). The ability of stem cells to preserve proliferation while progressively generating more differentiated progeny is achieved through asymmetric divisions, a key feature of neuroblasts (the stem cells of the central nervous system in *Drosophila*). Some aspects of asymmetric division are conserved between *Drosophila* and humans and involve the segregation of fate determinants, whereby molecules for sustaining proliferation are segregated apically to be maintained in the neuroblast (NB), while molecular cues guiding differentiation are positioned basally, to be segregated into the daughter cell for its differentiation (Homem and Knoblich 2012). Disruption in the cell type-specific expression of cell fate determinants can lead to uncontrolled proliferation and brain tumors or insufficient neural populations. During larval development, NBs are found throughout the larval brain lobes and ventral nerve cord. They are identified by their patterns of division, genetic markings, and positions within the brain. NB progeny are distinguished by their positions and genetic markers. Type I neuroblasts express both Deadpan and Asense and are found within the brain lobes and ventral nerve cord where they asymmetrically divide to self-renew and generate a more differentiated, Asense positive, ganglion mother cell (GMC). Type II neuroblasts are Deadpan positive and Asense negative (Supplemental Fig. 1C). They asymmetrically divide to generate an intermediate neural progenitor (INP). Once INPs mature, they become Asense and Deadpan positive and they themselves asymmetrically divide to generate a GMC which, similar to GMCs in the type I lineage, will symmetrically divide to generate two neurons or glia (Boone and Doe 2008; Bello et al. 2008; Bowman et al. 2008). The INP transit-amplifying pattern of divisions in type II neuroblast populations is similar to that found in the human cortex and leads to the generation of a much greater number of neurons, approximately 600 neurons by the type II lineage as opposed to around 100 neurons generated by the type I neuroblast lineage. The *Drosophila* larval nervous system provides a genetically tractable model to ask how ion channels influence cells in various states of proliferative potential and differentiation.

Previously, our lab has used *Drosophila* to characterize a role for the voltage-gated K+ channel ether-a-go-go (EAG) in tumor development (Huang et al. 2012; 2015). In this study, we examine how the voltage-gated sodium channel (VGSC) governs aspects of neural development and tumor proliferation. Overexpression of the pore-forming α subunit of VGSCs has been found in various cancers including breast cancer, cervical cancer, colon cancer, glioma, leukemia, lung cancer, lymphoma, melanoma, mesothelioma, neuroblastoma, ovarian cancer and prostate cancer (Patel and Brackenbury 2015; Fraser et al. 2005; Yang et al. 2012; Anderson et al. 2003; Abdul and Hoosein 2002; Xia et al. 2016). These studies implicate VGSCs in various aspects of cancer pathogenesis including migration, metastasis, invasion and proliferation. However, whether VGSC might also influence aspects of proliferation during normal development is not clear. There are 9 genes for pore-forming VGSC α subunits in the genome of mammals such as mouse and human. The *Drosophila* genome encodes a sole VGSC α subunit, Paralytic (*para*), rendering *Drosophila* a more straightforward model to investigate the *in vivo* role of VGSCs in stem cell and tumor proliferation. We have ided a role for *para* in regulating important aspects of neural progenitor proliferation in *Drosophila* larvae. Furthermore, we found that reduction of Para is sufficient to suppress brain tumor models driven by DeadpanOE (ectopic overexpression) (Zhu et al. 2012; Huang et al. 2015), activated Notch (Song et al. 2011, Zhu et al. 2012), or knockdown of Brat (Bowman:2008ib), indicating that Para may act downstream of genetic cascades that are known to regulate important aspects of proliferation and differentiation.

## Results

### Reduction or loss of Para compromised proliferation of type I and type II neuroblast lineages

To examine the role of Para in brain development, we used RNAi to knock down Para in the type I and type II neuroblast lineages using *inscuteable-Gal4* (*insc-Gal4*) (Supplemental Fig. 1A-C). We found that knockdown of Para resulted in volume reduction of brain lobes but not ventral nerve cord (Fig. 1A-C). To assess the involvement of Para in Type I and II neuroblast lineages, we generated a null allele of *para* through FLP recombinase of FRT insertion sites flanking the *para* gene region (method described in (Parks et al. 2004), Supplemental Fig. S2A-E). As Para represents the sole voltage-gated sodium channel in flies, its loss results in lethality (Broadie and Bate 1993). With MARCM (mosaic analysis with a repressible cell marker), we generated homozygous null clones within an otherwise heterozygous and viable animal (Lee et al. 1999).

**Fig. 1.**
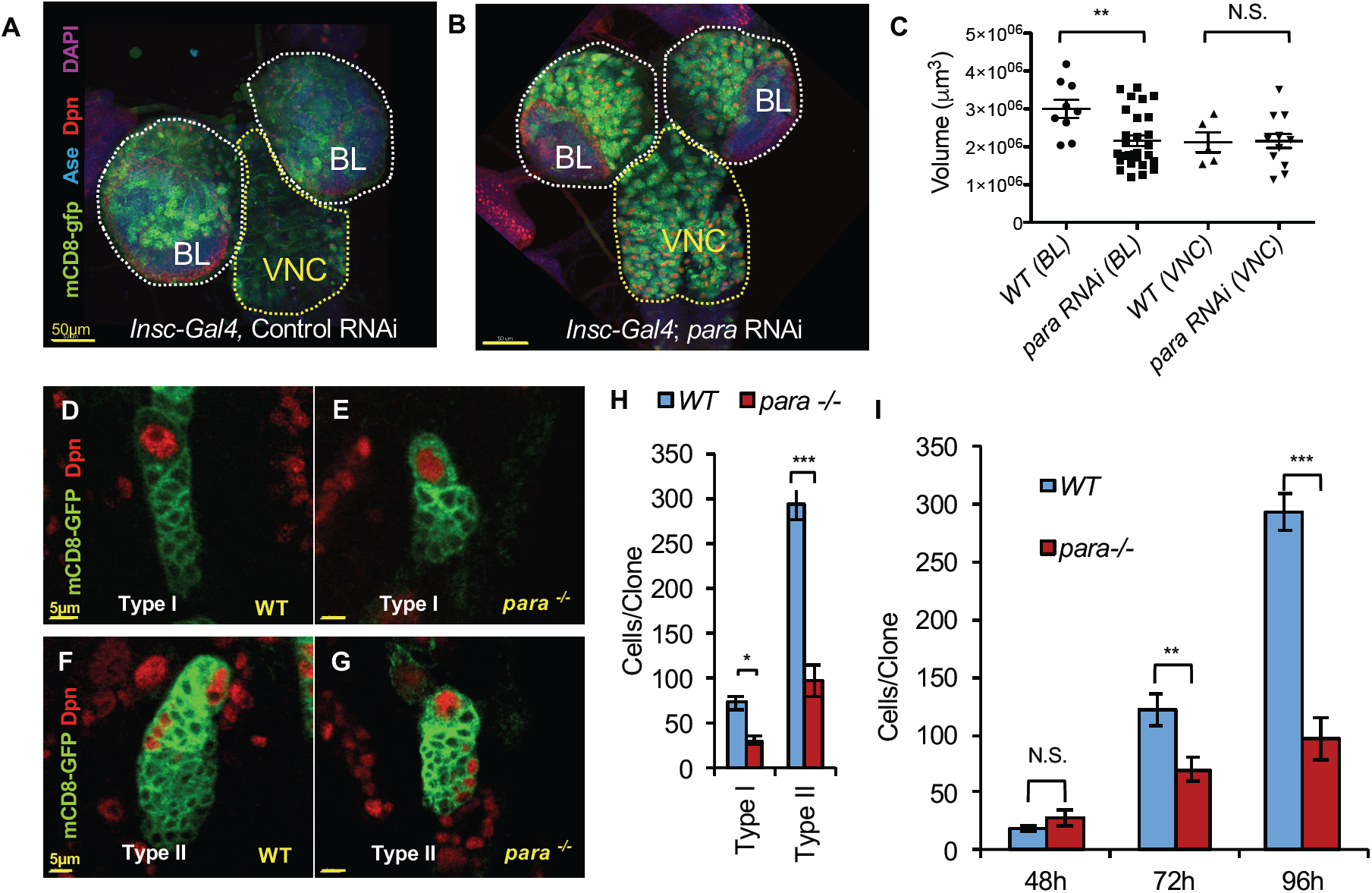
Reduction or loss of Para compromised proliferation of type I and type II neuroblast lineages. (A) Representative image of wildtype (WT) larval central nervous system composed of 2 brain lobes (BL, white) and a ventral nerve cord (VNC, yellow) Scale bar = 50µm. (B) RNAi knockdown of *para* in neural progenitors with *inscuteable -GAL-4* (*insc-GAL4*) in WT brains significantly reduced BL volume but not VNC volume. Scale Bar = 50µm. (C) BL: WT N=9, *para* RNAi N=28, VNC: WT N=5, *para* RNAi N=13 (D-E) Representative images of type I neuroblast WT and *para*^*-/-*^ MARCM (Mosaic Analysis with a Repressible Cell Marker) clones driven by enhancer trap FLP recombinase 200c (ET-FLP 200C) with *insc-GAL4* at 96 hours (h) after larval hatching (ALH) Scale bar = 5µm. (F-G) Representative images of type II WT and *para*^*-/-*^ ET-FLP 200c MARCM clones. (H) *para*^*-/-*^ type I and type II neuroblast MARCM clones had fewer cells per clone than WT. (type I: WT N=14, para-/- N=9, type II: WT N=20, *para*^*-/-*^ N=26). Two-tailed t-test, p<0.05. (I) Type II (ET-FLP200c) *para*^*-/-*^ MARCM clones displayed similar cell numbers at 48 hours after larval hatching (ALH), but at later time points, possessed progressively fewer cells per clone, compared to WT. (48h ALH: WT N=19 *para*^*-/-*^ N=5, 72h ALH: WT N=15 *para*^*-/-*^ N=18, 96h ALH: WT N=20, *para*^*-/-*^ N=26) Scale bar is 5µm.

*para*^-/-^ clones were marked by membrane bound mCD8GFP. We found that, compared to wildtype, *para*^-/-^ MARCM clones had fewer cells per clone in both type I and type II neuroblast lineages (Fig. 1D-H; Supplemental Fig. S1D). MARCM only removed *para* within the clone, which suggested that Para acts cell autonomously in neuroblast lineage development. Indeed, cell-autonomous expression of *para* cDNA within the type I (Fig. 2H-K) or type II (Fig. 2A-D) neuroblast lineage was sufficient to rescue cell number in *para* null clones at 72h after larval hatching (ALH), as well as at 96h ALH (Supplemental Fig. S3). By examining cellular subtypes within *para*^-/-^ MARCM clones, we found that loss of *para* significantly reduced the numbers of INPs, GMCs and neurons within type II clones, and this loss was rescued by *para* cDNA expression (Fig. 2A’-C’,E-G). Similarly, loss of *para* significantly reduced the numbers of GMCs and neurons within type I MARCM clones, and this loss was rescued by *para* cDNA expression (Fig. 2H’-J’,L-M). Notably, the role of Para in neuroblast lineages may be more specific to central brain lineages as there was no significant difference found for medulla neuroblast lineages within the optic lobe (Supplemental Fig. S4A-C). As the cellular deficit was stronger within the type II lineage and progressively worsened over time (Fig. 1I), we decided to focus our studies on the role of Para in the type II neuroblast lineage.

**Fig. 2.**
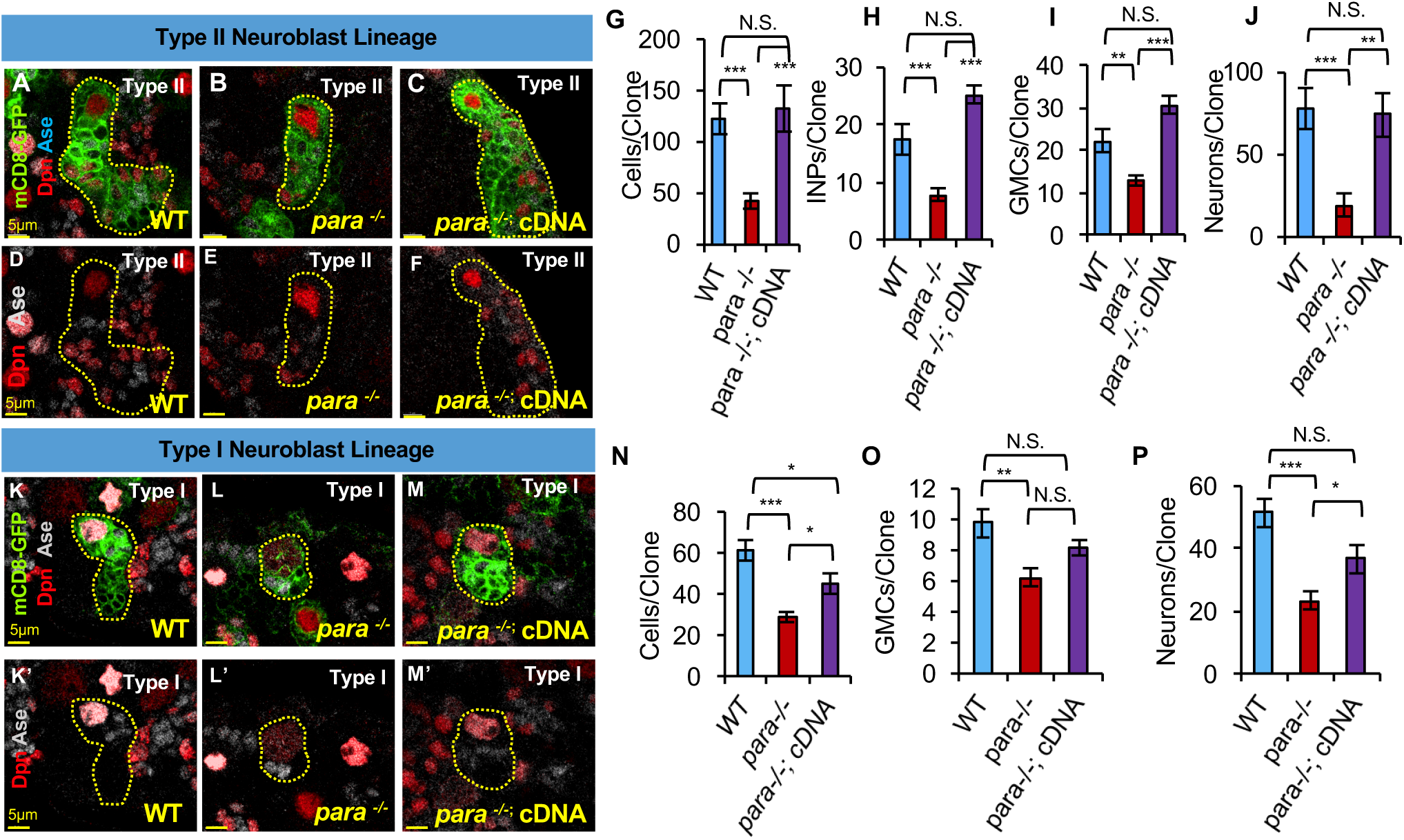
Cell loss in type I and II *para*^*-/-*^ MARCM clones is rescued by cell autonomous expression of para cDNA. (A-C) Representative images of type II MARCM clones (driven by a heat shock (HS) FLP recombinase with insc-GAL4). Scale bar = 5µm. (G) 72h ALH *para*^*-/-*^ reduction in clones per MARCM clone compared to WT is rescued by cell autonomous expression of a *para* cDNA. One-way ANOVA with Bonferonni multiple comparison test, *** p<0.0001 WT N=14 *para*^*-/-*^ N=19 *para*^*-/-*^; *para* cDNA N=11 (D-F) Representative images of of hs-FLP 72h ALH type II MARCM clones (at higher magnification and maximum projection to show progeny). INPs marked by Deadpan (Dpn^+^) in red and Asense (Ase^+^) in grey, GMCs marked by Ase^+^ alone. Scale bar = 4µm. (H-J) Type II MARCM clone cellular subtypes (Intermediate neural progenitors (INP), ganglion mother cells (GMCs), neurons) are reduced compared to WT and this loss is rescued by cell autonomous expression of a *para* cDNA. Clones are 72h ALH, P<0.05 (H) INPs: WT N=14, *para*^*-/-*^ N=19, cDNA rescue N=8 One-way ANOVA with Bonferonni multiple comparison test, p<0.001 (I) GMCs: WT N=14, *para*^*-/-*^ N=18, cDNA rescue N=5, One-way ANOVA with Bonferonni multiple comparison test, p<0.01 (J) Neurons: WT N=14, *para*^*-/-*^ N=18, cDNA rescue N=5. One-way ANOVA with Bonferonni multiple comparison test, p<0.01 (K-M) Representative images of type I MARCM clones (driven by a heat shock (HS) FLP recombinase with insc-GAL4). Scale bar = 5µm. (G) *para*^*-/-*^ MARCM clone cell number deficit is rescued by cell autonomous expression of a para cDNA, time = 72h ALH. One-way ANOVA with Bonferonni multiple comparison test, *** p<0.0001 (72h ALH: WT N=17 *para*^*-/-*^ N=20 *para*^*-/-*^; *para* cDNA N=14). (K’-M’) Representative images of of hs-FLP 72h ALH type I MARCM clones. Scale bar = 5µm. (N-P) Type I MARCM clone cellular subtypes GMCs, neurons) are reduced compared to WT and this loss is rescued by cell autonomous expression of a *para* cDNA. Clones are 72h ALH, P<0.05 (H) INPs: WT N=14, *para*^*-/-*^ N=19, cDNA rescue N=8 One-way ANOVA with Bonferonni multiple comparison test, p<0.05 (I) GMCs: WT N=17, *para*^*-/-*^, cDNA rescue N=14, One-way ANOVA with Bonferonni multiple comparison test, p<0.01 (J) Neurons: WT N=17, *para*^*-/-*^ N=14, cDNA rescue N=5. One-way ANOVA with Bonferonni multiple comparison test, p<0.05

### Type II *para*^-/-^ MARCM clones displayed a reduced rate of cellular accumulation

Loss of *para* led to a reduction of cell numbers per MARCM clone and the rescue data suggested that *para* acts cell autonomously. To characterize cell loss in type II *para*^-/-^ MARCM clones, we examined the incorporation of EdU (5-ethynyl-2-deoxyuridine), a thymidine analog that is incorporated into DNA of dividing cells and labels newly generated progeny. Larvae were fed EdU for 4 hours. After this time, we dissected a fraction of these larvae immediately off Edu (T = 0 hours (h)), so that most dividing cells (NB, INPs, GMCs) would be labeled. At a later timepoint, 12h off EdU, EdU would increasingly label post mitotic cells and be diluted from proliferating cells as they continue to divide (Fig. 3A,D-E’). In wildtype type II MARCM clones, cell number increased over 12 hours and, as expected, the number of EdU+ cells increased as proliferation diluted EdU into progeny (Figure 3B-C, F, F’). *para*^-/-^ MARCM clones displayed fewer cells at T = 0h off EdU compared to wildtype (Fig. 3B). However, in contrast to WT clones, the number of cells, as well as EdU labeled cells, was not significantly different between T = 0h and T = 12h in *para*^-/-^ clones (Fig. 3B,C, E, E’, G, G’). This suggests that type II *para*^-/-^ MARCM progenitors either proliferated more slowly or have increased cell death. As the number of EdU incorporated cells was comparable between *para*^-/-^ and WT at T = 0h, it may indicate that a similar fraction of mitotically active cells were capable of taking up EdU during the 4 hour time frame and that cell cycle speed may not be a major contributor to the differences in cell number at T = 12h (Fig. 3C).

**Fig. 3.**
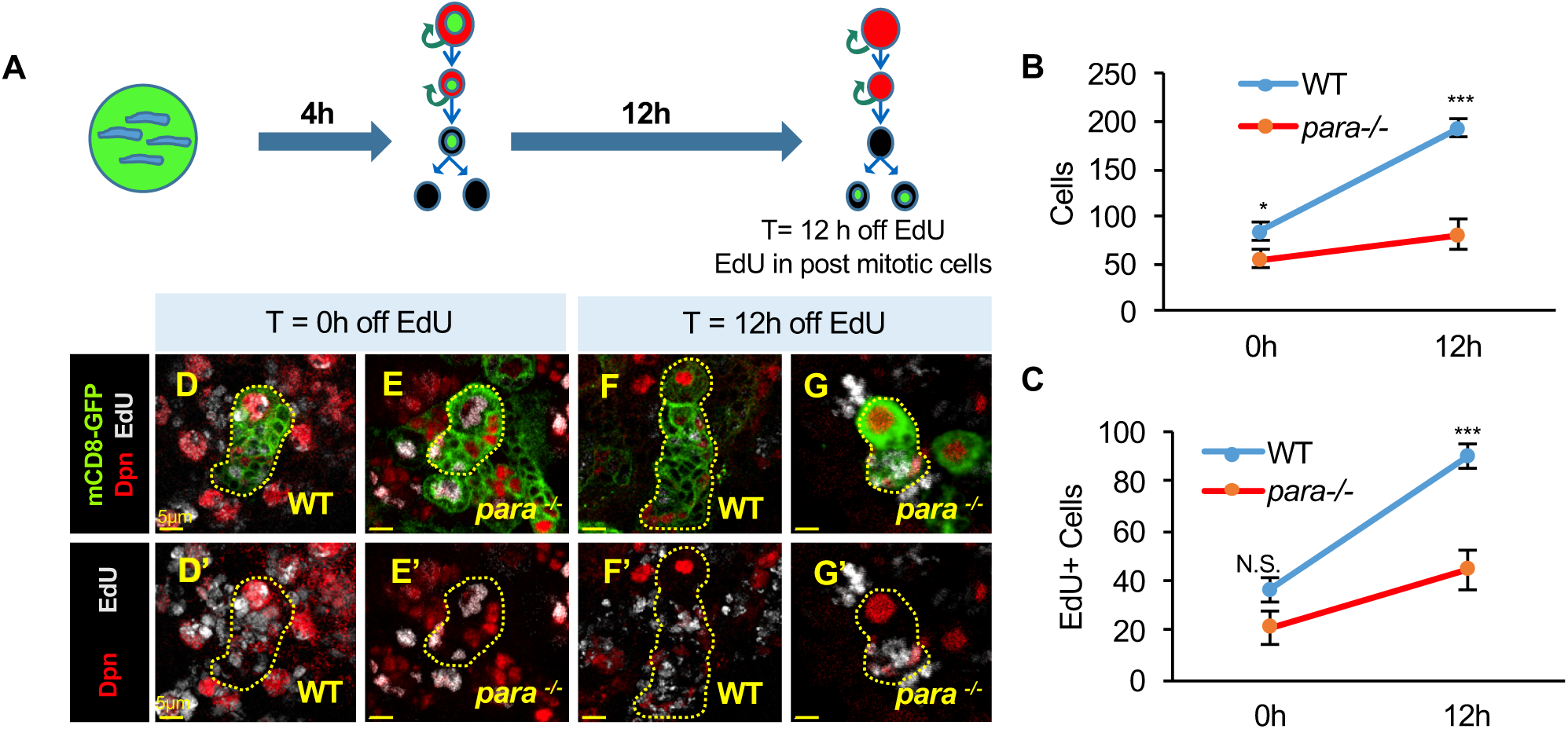
Type II para-/- MARCM clones displayed a reduced rate of cellular accumulation. (A) Schematic of EdU (5-ethynyl-2-deoxyuridine) protocol. Larvae staged at 72h (hours) after larval hatching (ALH), are fed media mixed with EdU. (D-E’) Representative image of wildtype (WT) MARCM clone at time = 0h off EdU, scale bar = 5µm. Dissections occurred immediately following EdU feeding period, indicated as time = 0h off EdU. At this time point, EdU (in grey) colocalizes with mitotically active neuroblasts (NBs) and intermediate progenitors (INPs) marked by Deadpan (Dpn) staining in red, as well as in Dpnganglion mother cells (GMCs). Membrane bound mCD8-GFP labels MARCM clones in green. (F-G’) Representative image of WT MARCM clone at 12h off EdU, scale bar = 5µm. At 12h off EdU, mitotically active cells increased numbers of EdU labeled cells within MARCM clones, as EdU was diluted from mitotically active cells into new born progenitors and post mitotic neurons. At At 12h off EdU, fewer Dpn^+^ cells colocalized with EdU (F-G’). (B) At T=0h off EdU, type II *para*^*-/-*^ MARCM clones display fewer cells than WT. (T=0h off EdU: WT N=7 *para*^*-/-*^ N=9, two-tailed T-test, P<0.05) At 12h off EdU, the difference in cell number, between WT and *para*^*-/-*^ MARCM clones, is even greater, as *para*^*-/-*^ type II MARCM clone cell numbers were not increased to the same extent as WT. (T=12h off EdU: WT N=8 *para*^*-/-*^ N=7, two-tailed T-test, P<0.05). (C) At t=0h off EdU, WT and *para*^*-/-*^ have incorporated similar numbers of EdU labeled cells, at T= 12h off EdU, the number of EdU_+_ cells in *para*^*-/-*^ type II MARCM clones was not increased to the same extent as WT. (T=12h off EdU: WT N=7 *para*^*-/-*^ N=7, two-tailed T-test, P<0.0001).

### Apoptosis is a major contributor to *para*^-/-^ type II lineage cell loss

To determine whether *para*^-/-^ cells undergo apoptosis, we made use of the baculovirus P35 to block apoptosis. During normal development, a significant proportion of type II neuroblast derived neurons die (Jiang and Reichert 2012). Consistent with this observation, we found that blocking cell death in WT type II lineages led to a slight increase in cell number (Fig. 4A, C-D). In contrast, blocking apoptosis in type II *para*^-/-^ MARCM clones dramatically increased the number of cells per clone (Fig. 4A,E-F). This increase in cell number of P35 expressing *para*^-/-^ clones rescued *para*^-/-^ cell number to a level similar to that of WT clones expressing P35 (Fig. 4A). These data suggested that the major driver of *para*^-/-^ cell loss was due to apoptosis. We confirmed these findings by assessing the cleaved caspase (cDCP-1) staining in type II lineages and found that RNAi knockdown of *para* reduced the number of cells per lineage and increased cDCP-1 labeled cells, which indicated increased cell death upon reduction of Para (Supplemental Fig. 5A-F). Interestingly, while WT INP numbers were unchanged compared to WT clones expressing P35, *para*^-/-^ clones expressing P35 had more INPs than those not expressing P35 (Fig. 4B,C’-F’). Consistent with this idea, knockdown of *para* in type II lineages led to an increase in Ase+ cDCP1+ labeled cells (Supplemental Fig. 5A’-C’, G), indicating that without Para INPs and possibly GMCs underwent cell death. To investigate how INPs might be lost to apoptosis in *para*^-/-^ clones, we used a cell cycle reporter (Supplemental Fig. S6A-C), and found an increased percentage of *para* null type II MARCM progeny (INPs and GMCs) in G2/M phase and a decreased percentage of cells in G1 as compared to WT (Supplemental Fig. S6E). As INPs and GMCs are much smaller than neuroblasts and thus more difficult to assess for DAPI condensed chromosomes, we stained for phospho histone 3 (pH3), a M-phase marker, and found that an increased proportion of INPs are pH3+ and likely in M-phase (Supplemental Fig. S6F-H’).

**Fig. 4.**
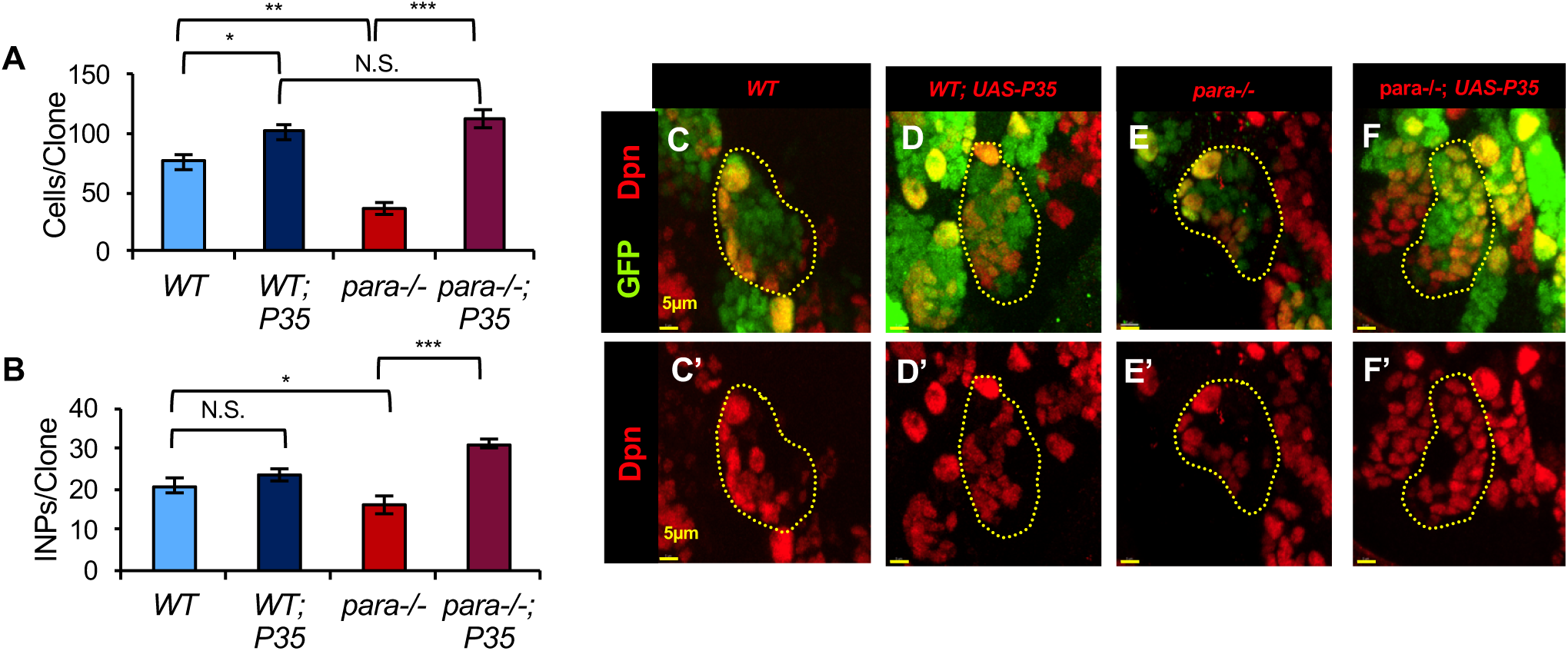
Apoptosis is a major contributor to *para*^*-/-*^ type II lineage cell loss. (**A**) Baculovirus P35 was expressed within clones to block apoptosis. Expression of P35 slightly increased the number of WT cells, which likely accounts for the fraction of wildtype (WT) cells which undergo apoptosis normally during development. Blocking apoptosis in *para*^*-/-*^ type II MARCM clones completely rescues the number of cells to WT + UAS-P35 levels, indicating that cells may cycle more slowly, but ultimately cell death is a major driver of cell number reduction in *para*^*-/-*^ MARCM clones. (One-way ANOVA with Tukey multiple comparison test, p<0.05. WT N=10 WT UAS-P35 N=9 para-/- N=6 *para*^*-/-*^; UAS-P35 N=8). (B) *para*^*-/-*^ MARCM clones have fewer intermediate progenitors (INPs) than WT. Blocking cell death with P35 expression does not change the number of INPs in WT MARCM clones indicating that other cells may be dying. Blocking cell death in *para*^*-/-*^ MARCM clones increased the number of INPs indicating that INPs are one of the cell populations sensitive to apoptosis in para-/- MARCM clones. (One-way ANOVA with Tukey multiple comparison test, p<0.05. WT N=11 WT P35 N=9 *para*^*-/-*^ N=9 *para*^*-/-*^ P35 N=8). (C-F’) Representative images of type II MARCM clones marked by GFP NLS (nuclear localized, green fluorescent protein) in green and stained for Dpn (Deadpan) in red to mark neuroblasts and INPs. Scale bar is 5µm.

Conceivably, loss of Para may slow the cell cycle so as to increase the fraction of INPs in M-phase, and in some cases resulting in cell cycle arrest that culminates in cell death. Loss of INPs would have cascading effects, diminishing the number of GMCs and neurons. There was also an enrichment of *para*^-/-^ type II neuroblasts in G2 phase (Supplemental Fig. S6A-D), suggesting that these cells cycle more slowly than wildtype. Additionally, as neural activity has been shown to be important for axon guidance and synaptic refinement (Casagrande and Condo 1988; Patel and Brackenbury 2015), and loss of Na_V_ 1.2 results in marked increases in neuronal apoptosis (Planells-Cases et al. 2000), a proportion of *para*^-/-^ neurons may be expected to die from apoptosis. While we did not see increased cDCP1 staining within neurons (Supplemental Fig. 5H), *insc-Gal4* was more strongly expressed in NBs and INPs. Moreover, when a more direct neuronal Gal4 was used (*elav-Gal4*), no progeny arose, indicating that knockdown of Para directly within neurons is lethal as has been previously reported (data not shown and Parker:2011el). Thus, we cannot exclude the possibility that neuronal cell death may occur with a greater reduction of Para or at other time points we did not investigate. These data indicated that loss of *para* activity within MARCM clones led to fewer cells due to apoptosis of neuroblast progeny which potentially arose from cell cycle arrest, and may have minor contributions from a slower cell cycle of neuroblasts and neural progenitors.

### Overexpression of Para increased cell number in type II NB lineages

Having found that loss of *para* reduced cell number in *Drosophila* neuroblast lineages, we examined the effect of increasing Para expression in an otherwise wild-type lineage. To this end, we overexpressed a wildtype *para* cDNA as well as a *para* gain of function allele, bang senseless (BSS), which is thought to cause hyperexcitability via a shift of fast inactivation towards more positive potentials (Parker et al. 2011). bang senseless was identified as a seizure prone mutant upon mechanical stimulation (Ganetzky and Wu 1982) and later attributed to the missense mutation L1699F within the *para* locus (Parker et al. 2011).The L1699 residue is highly conserved among mammalian VGSCs and lies within transmembrane segment S3 of the fourth homology domain of the Para protein. Overexpression of *para* cDNA in the type II neuroblast lineage, using *insc-Gal4* with *ase-Gal80*, increased the number of cells per lineage compared to wildtype at the same time point (Fig. 5A-D). There was no significant difference between WT *para* cDNA over-expression and bss cDNA overexpression (Fig. 5D). Notably, INP numbers were increased in *para* cDNA overexpressing lineages compared to WT, indicating that increased expression of Para may lead to faster proliferation of type II neuroblasts or longer lifespans of INPs (Fig. 5A’-C’, E). Consistent with the idea that INPs may be regulated by Para, altering *para* expression levels with RNAi knockdown (Supplemental Fig. 7A-F) or by overexpression of *para* cDNA in INPs alone was sufficient to decrease or increase INP number as well as the numbers of GMCs and neurons, respectively (Supplemental Fig. 7G-L). Thus *para* expression likely influences NBs as well as INPs.

**Fig. 5.**
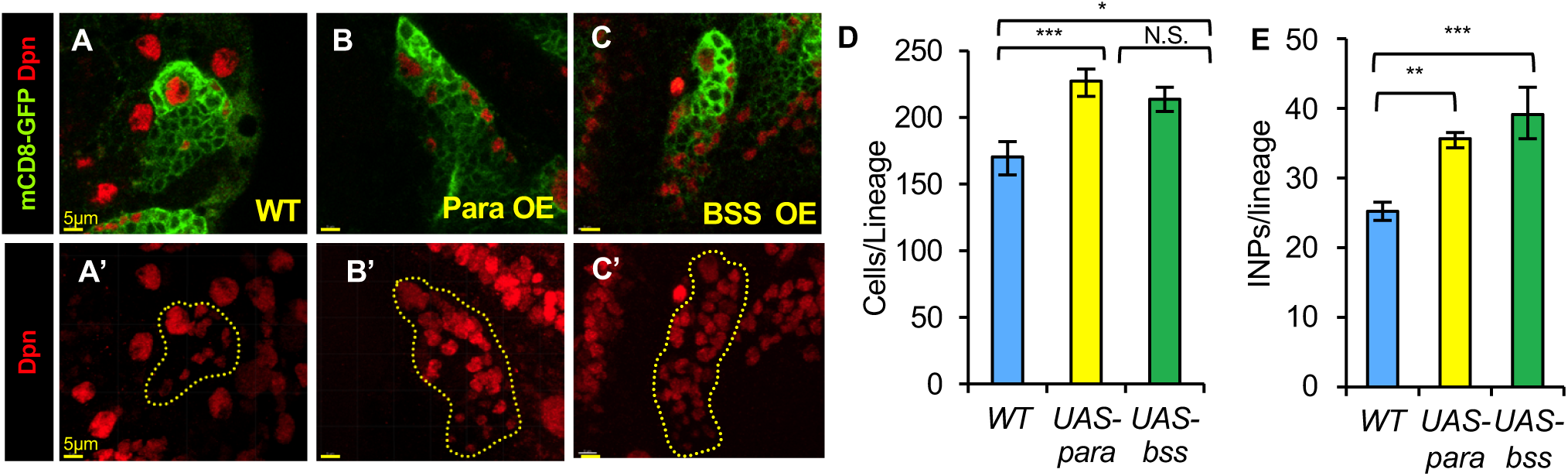
Overexpression of para cDNA increased cell number in WT type II lineage. (A-D) Overexpression of wildtype (WT) cDNA labeled Para OE (Para Over Expression) or overactive para seizure-associated allele, bang senseless (BSS OE), driven by incuteable-GAL4; asense-GAL80, increased the number of cells per clone compared to WT type II lineages (marked by membrane bound mCD8-GFP in green) at 96h ALH (after larval hatching) (D). WT N=13, Para WT cDNA N=15 WT BSS cDNA N=12. One-way ANOVA with Bonferonni multiple comparison test, P<0.05. (A-C) Representative images of WT and para cDNA overexpression. Scale bar = 5µm. (A’-C’) Maximum projections of WT or para cDNA overexpression in type II cellular lineage. Para OE and BSS OE showed more INPs marked by Deadpan (Dpn) in red than WT, scale bar = 5µm (E) WT N=18, Para WT cDNA N=12 WT BSS cDNA N=5. One-way ANOVA with Bonferonni multiple comparison test, P<0.01.

### Reduction of Para suppressed brain tumor size

Having found that the abundance of Para influences proliferation in neuroblast lineages, we asked whether loss of Para can influence overproliferation within a brain tumor. Deadpan (Dpn) is a bHLH (basic-helix-loop-helix) transcriptional repressor expressed in neuroblasts of embryonic and larval brains, as well as intermediate neural progenitors (Fig. 6A-A”) (Younger-Shepherd et al. 1992; Bier et al. 1992). Over-expression of Dpn in the type I and type II neuroblast lineage, driven by the *insc-Gal4*, led to brain tumor formation in *Drosophila* larval brains as a result of overproliferation (Fig. 6B-B”,D-D”,F-G) (Zhu et al. 2012). RNAi knockdown of *para* within a DpnOE tumor resulted in a reduction in size of both the brain lobes and ventral nerve cord (Fig. 6C-C”,E-E”,F-G). This reduction of tumor mass within brain lobes and ventral nerve cord indicates that knockdown of Para may influence tumor cells derived from both type I and II neuroblasts. A recent study examined the crystal structure of Brat, a translational repressor, in complex with RNAs. Brat is involved in direct differentiation of neuronal stem cells by suppressing self-renewal factors, and it binds known proliferation factors including *chinmo, dpn, klu, staufen(stau)*, and *par-6* mRNAs (Loedige et al. 2015). An unexpected finding was that Brat was also found in complex with *para* mRNA, suggesting a potential influence of the sole *Drosophila* VGSC in promoting stemness. To ask whether Para may have a more generalized role in proliferation, we generated type II neuroblast derived tumors with genes involved in developmental pathways known to influence neuroblast lineage self-renewal and differentiation. Overexpressing Dpn (DpnOE)Zhu:2012eh, overexpressing activated Notch (NIC) Zhu:2012eh, Song:2011eb which is important for stem cell maintenance, or knockdown of Brat (a translational repressor of stem-cell promoting genes)Bowman:2008ib with *insc-Gal4* with *ase-Gal80* was sufficient to generate type II brain tumors (Figure 6H-J). Introducing a hypomorphic *para* mutant, *paraTS1*, in these type II neuroblast derived tumors, led to a profound reduction in tumor size (Fig. 6K-N). Together these data indicated that *para* promoted brain tumor growth derived from both type I and II lineages, as knockdown of *para* with RNAi or use of a hypomorphic *para* mutant allele, both suppressed brain tumor mass. Our findings that the *para*^*TS1*^mutation is capable of suppressing tumors generated by manipulating Dpn, Notch and Brat signaling suggest that it regulates proliferation downstream of important developmental cascades (Supplemental Fig. S8D).

**Fig. 6.**
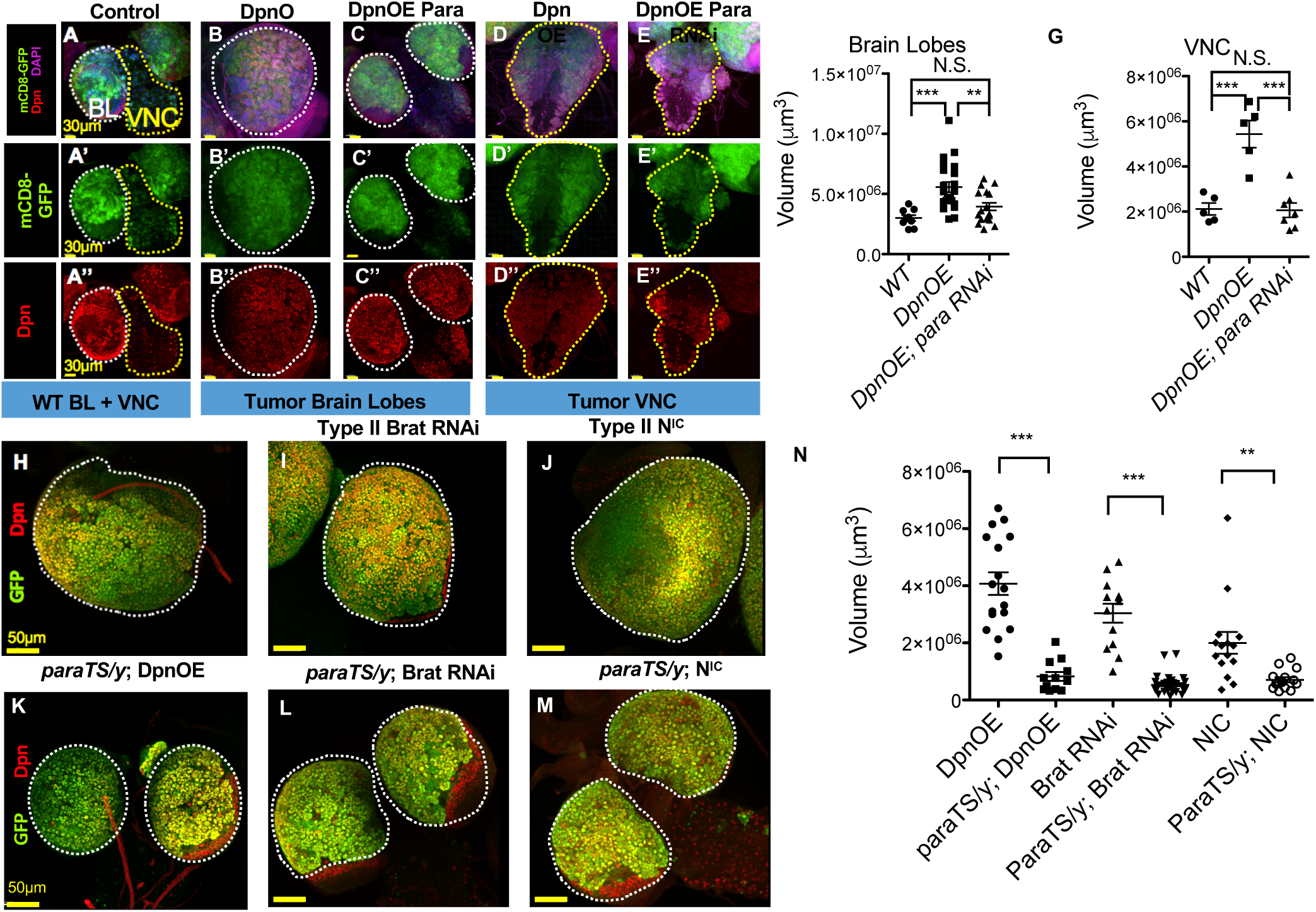
Reduction of Para expression suppressed Drosophila brain tumors. (A-A”) Representative image of WT CNS, composed of 2 brain lobes (BL) and a single ventral nerve cord (VNC) Scale bar = 30µm. (B-B”, D-D”) Ectopic overexpression of Dpn, driven by insc-GAL4, led to expansion of Dpn^+^ cells (marked in red (B”)) and generation of a brain tumor with enlarged brain lobes (B-B” and F) and ventral nerve cord (VNC) (D-D” and G). Scale bar = 30µm. (C-C”, E-E”) *para* RNAi knockdown in DpnOE brain tumor resulted in reduced BL (F) and VNC volume (G). One-way Anova with Tukey multiple comparison test, P value <0.05 (BL: DpnOE, N=21, DpnOE *para* RNAi, N=17) and (VNC: DpnOE, N=5, DpnOE *para* RNAi, N=8) (H-J) Representative image of type II brain tumors driven by *insc-GAL4* with *asense-Gal80* to limit expression to only the type II, brain lobe restricted (and nuclear localized green fluorescent protein (GFP) for volume measurement) (H) DpnOE brain tumor, generated by Dpn overexpression (I) Brat brain tumor, generated by *brat* RNAi knockdown, Notch (NIC) brain tumor generated by overexpression of activated Notch (J-L) Representative image of hypomorphic *para*^*TS1*^ mutants reduced type II brain tumors. Scale bar = 50µm (M) One-way ANOVA with Dunnett’s multiple comparison test, *** p<0.0001 (BL: DpnOE, N=17, *para*^*TS1*^*/y*, N=12, Brat, N=13, *para*^*TS1*^*/y*; Brat, N=37, NIC, N=17, *para*^*TS1*^*/y*; NIC, N=12).

### Para channel activity is important for its role in type II neuroblast lineage development

We next asked whether the involvement of Para in development depended on its function as a conduit for ion flow. Para is a pore forming α subunit member of the VGSC family; it contains four homologous domains (DI-DIV) each with 6 transmembrane segments (Fig. 7A) (Catterall 2000). One α subunit forms a functional channel, but the channel activity and location can be influenced by subunits (Payandeh et al. 2011). VGSCs are closed at the resting membrane potential. Upon membrane depolarization, VGSCs are activated through outward movement of the S4 voltage sensors. After a few milliseconds, VGSCs inactivate through an inactivation gate composed of the intracellular loop connecting domains III and IV (Catterall 2000). To address whether the Para ion channel activity was important in development, we examined a lethal point mutant *para*V1401E (Yamamoto et al. 2013). Valine 1401 lies within the intracellular loop between S4 and S5 of the third homology domain (Fig. 7A), where the change from the hydrophobic Valine to the hydrophilic residue Glutamate may result in misfolding or interfere with voltage gating. In addition, residues within this intracellular loop IIIS4-S5 are thought to stabilize the inactivation gate, and mutations in this region appear to impair inactivation (Smith and Goldin 1997; Catterall 2000). We found that homozygous MARCM clones of *para*V1401E displayed fewer cells per clone and phenocopied *para*^-/-^ MARCM clones (Fig. 7B). As this mutant allele had not been electrophysiologically characterized, we performed whole cell patch clamp recordings of WT *para* and *para*V1401E mutant channels heterologously expressed in Xenopus oocytes. The tipE subunit was co-injected in a 1:1 molar ratio to stabilize expression (Feng et al. 1995; Warmke et al. 1997). Oocytes were recorded using two-electrode voltage clamp to obtain current-voltage (I-V) relationships from a holding potential of −80 mV. Wild-type Para channels demonstrated a mean peak current amplitude at 0 mV of −0.87 +/− 0.11 µA (Fig. 7C; Supplemental Fig. S9A). The V1401E mutant channels displayed a significant reduction in current amplitude, with a mean peak of −0.11 +/− 0.01 µA (Fig. 7C; Supplemental Fig. S9B). In order to probe whether Na+-flow or some other property of Para protein was responsible for the phenotypes, we designed a “pore dead” construct based on homology with Na_V_ 1.2. To this end we performed a full protein sequence alignment (ClustalD) of Para with rat Na_V_ 1.2 (NP 036779.1) and replaced a single aspartate residue, D388, with as*para*gine. This residue contributes to the conserved ion conduction pore among Nav family members (Fig. 7A), and mutation to as*para*gine results in reduced sodium conductance while preserving surface expression of the channel (Pusch et al. 1991). When expressed in oocytes, D388N mutant channels yielded currents barely distinguishable from background with a mean peak current amplitude of −0.07 +/− 0.01 µA (Fig. 7C; Supplemental Fig. S9C). These results suggested that *para*V1401E had impaired channel function and, as it phenocopied complete loss of *para*, that channel function was likely important for the contribution of *para* to the development of neuroblast lineages. To further examine the importance of channel function, we asked whether the non-conducting D388N mutant channel is capable of rescuing *para*^-/-^ MARCM clones and whether its overexpression in the type II wildtype lineage would lead to increased numbers of cells, as was the case for WT *para* cDNA. *para*D388N cDNA was not able to rescue *para*^-/-^ MARCM clone cell numbers (Fig. 7D,F-I), nor increase the number of cells per type II neuroblast lineage (Fig. 7E,J-L), suggesting that the ability of Para to conduct Na+ ions is important for its role in neuroblast development. Together these data suggest that the Para voltage-gated sodium channel activity is important for its role in neuroblast development.

**Fig. 7.**
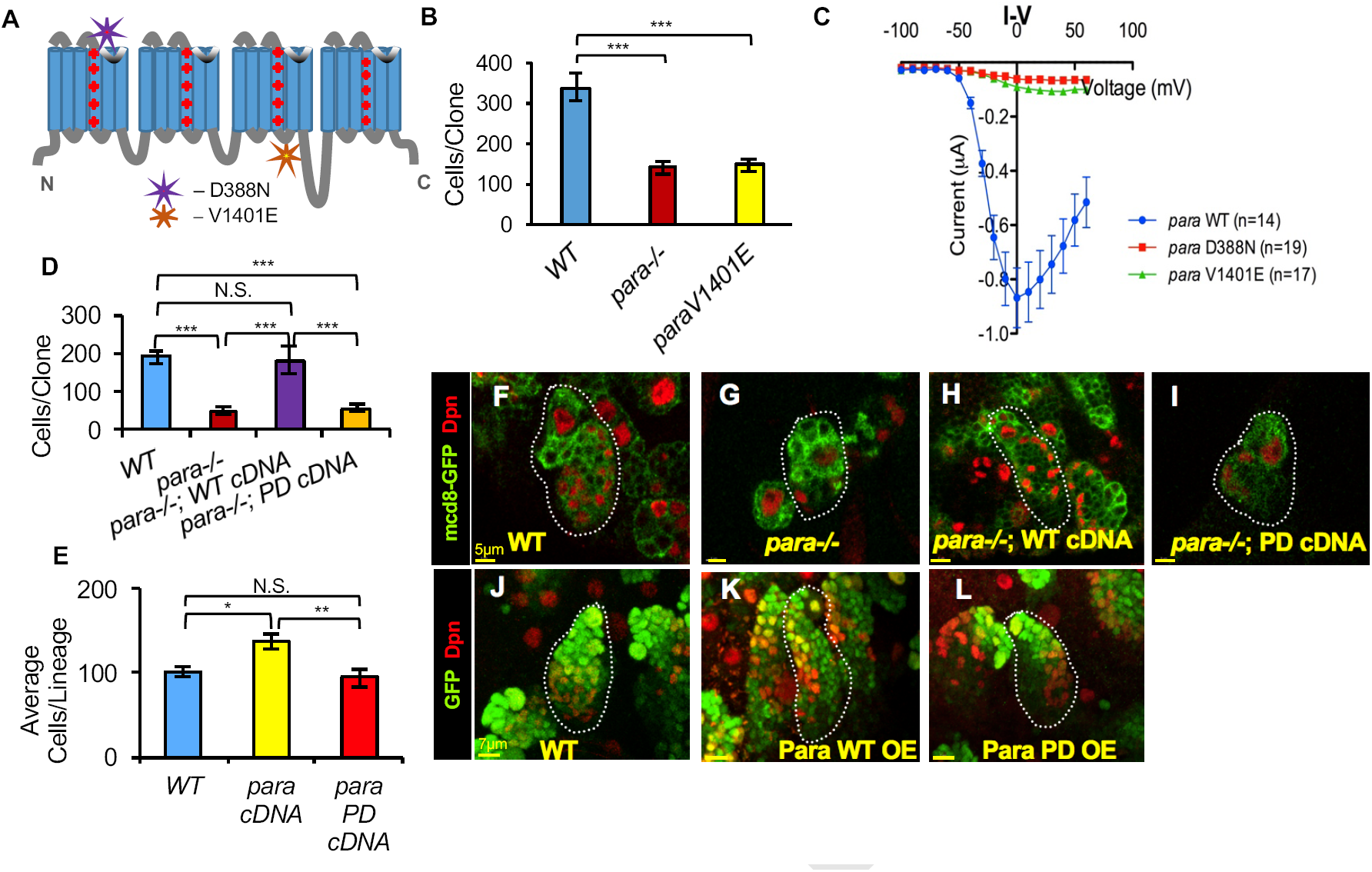
Para channel activity is important for its role in type II neuroblast lineage development. (A) Cartoon of Para protein structure showing relative location of missense point mutants. (B) *para*^*V1401E*^ phenocopied *para*^*-/-*^ type II MARCM clones, displaying fewer cells per clone compared to WT, at 96 hours after larval hatching (ALH). (One-way ANOVA with Tukey’s multiple comparison test, *** p<0.0001. WT N=19 *para*^*-/-*^ N=20 *para*^*V1401E*^ N=29). (C) Whole-cell Na+ currents in cRNA-injected oocytes were measured using a two-microelectrode voltage clamp to examine the current–voltage relationship (I-V curves) of WT versus V1401E and a D388N (aspartate to asparagine) mutation of a mammalian conserved residue, in the pore region, previously shown to be non-conducting (C). WT para currents are inward rectifying while V1401E and a D388N display little to no current WT N=14 V1401E N=16 D388N N=19. (D) Unlike WT *para* cDNA, para D388N cDNA was unable to rescue type II *para*^*-/-*^ cell deficit, indicating WT channel function is necessary for Para’s role in proliferation. (One-way ANOVA with Tukey’s multiple comparison test, *** p<0.0001 WT N=6 *para*^*-/-*^ N=8 *para*^*-/-*^ WT cDNA N=5 para-/- D388N cDNA N=5). (F-I) Representative images of type II MARCM clones marked by membrane bound mCD8-GFP. Scale bar=5µm. (E) Unlike WT *para* cDNA, overexpression of *para*^*D388N*^ mutant cDNA did not increase numbers of type II cells. (One-way ANOVA with Tukey’s multiple comparison test, *** p<0.05 WT N=6 WT cDNA OE N=7 D388N cDNA OE N=9). (J-L) Representative images of type II MARCM clones marked by nuclear localized GFP. Scale bar =7µm.

## Discussion

VGSCs play important roles during neural development where neural activity is important for axon guidance and synaptic refinement (Subramanian et al. 2012; Casagrande and Condo 1988). VGSC activity within the nervous system is essential, as evident from the fact that its absence leads to lethality in a number of mouse models (Harris and Pollard 1986; Planells-Cases et al. 2000; Yu et al. 2006). In addition, mutations in VGSC are associated with childhood diseases including autism spectrum disorders, epilepsy, mental retardation, among others, highlighting the necessity for understanding the role of VGSC during development (Eijkelkamp et al. 2012). While the function of VGSC in excitable tissue is well established, what they might be doing in non-excitable cells is not well understood. Recent work has begun to uncover functions of VGSC in development of non-excitable cells. One such study found that NaV1.3 channels (encoded by SCN3A) are expressed in radial glial cells and that they are important for migration during cortex development (Smith et al. 2018). A large scale clinical study finds that 25% of individuals with SCN2A mutations presented with microcephaly (Stessman et al. 2017), suggesting that VGSC may influence neuronal progenitor proliferation. Additionally, VGSCs are found to be critically important for cardiac progenitor development in zebrafish (Chopra et al. 2010). Taken together, these studies indicate that VGSCs play a role in progenitors during development, although the nature of their contributions is unclear.

In this study, we identified a role for the VGSC *para* in neural development. To our knowledge, our studies are the first to identify a role for VGSC in neural progenitor proliferation. We found that loss of *para* reduced total cell number in the type I and type II central neuroblast lineages compared to control. These effects are cell autonomous as MARCM clones homozygous for the *para* mutation displayed the phenotypes, which could be rescued by cDNA expression within these neuroblast lineages. The stronger phenotype observed in the type II lineage may reflect a role for Para in specific transit-amplifying cell population, INPs, which are only present in the type II lineage. Furthermore, loss of Para led to increased apoptosis that may arise from a failure of cell cycle progression. Interestingly, research in other systems has found that depolarization is essential for G2 to M progression (Blackiston et al. 2014). Perhaps Para promotes cell cycle progression in *Drosophila* progenitors through its depolarizing activity (Supplemental Fig. S6B). Generally, proliferative cells display more depolarized resting membrane potentials, as compared to differentiated cells, like neurons (Yang et al. 2012). In this context, Para may primarily exist in the inactivated state and a persistent sodium current, which is typically a few percent of the transient current, may predominate. In addition, VGSCs are highly subjected to RNA editing and alternative splicing. Some of these variants result in window currents and persistent currents, like that found in embryonic variants of *para* (Lin et al. 2009). Para could, therefore, contribute to a depolarized state that may be important for proliferation. Alternatively, *para* could be temporally upregulated during specific states of the cell cycle to activate down-stream molecular cascades. Depolarization has been shown to influence a number of aspects of cell cycle regulation, including changes in gene expression, calcium signaling, protein localization and phosphorylation states (McLaughlin and Levin 2018; Abdul Kadir et al. 2018). In addition, the contribution of VGSC to axon growth cones and metastasis and migration indicates a role for depolarization in cytoskeletal rearrangement (Patel and Brackenbury 2015). The cytoskeleton contributes to many aspects of cell division including subcellular localization of cell fate determinants during asymmetric cell division, volume changes, nuclear envelope breakdown and chromosome segregation during M-phase (Homem and Knoblich 2012; Hutterer et al. 2004; Betschinger and Knoblich 2004). If *para* activity were to influence any of these processes, dysregulation of cell cycle progression would be expected.

VGSCs drive membrane depolarization as they selectively permit permeation of sodium ions thereby driving the membrane potential towards the sodium equilibrium potential. Studies on regeneration in Xenopus laevis reveal that Na_V_ 1.2 mediated Na+ influx, rather than depolarization, is required for initiating regeneration following tail amputation (Tseng et al. 2010). This Na+ signaling, as a result of amputation, acts through salt inducible kinase (SIK) to drive proliferation, outgrowth and morphogenesis by activating Notch, Wnt, BMP and FGF signaling to complete regeneration (Tseng et al. 2010). Moreover, a number of recent studies in *Drosophila* find that sodium permeable ion channels promote proliferation. These include the epithelial sodium channel ENaC whose overexpression leads to overproliferation of gut stem cells (Kim et al. 2017), cation permeable TRPA1 channels that are upregulated in the gut in response to increased ROS so as to induce proliferation (Xu et al. 2017), and cation permeable Piezo that is important for proliferation in the adult fly midgut (He et al. 2018) as well as glioma proliferation in multiple fly glioma models (Chen et al. 2018). These studies highlight the potential role of Na+ currents as drivers of proliferation.

The effect of cell loss in type II *para*^-/-^ MARCM clones is progressive, with more severe loss at later time points. Neuroblasts and INPs undergo successive asymmetric divisions and these divisions act as a clock changing the transcriptional profile of these cells from early stages to late stages (Ren et al. 2017; Bayraktar and Doe 2013; Kohwi and Doe 2013; Li et al. 2013). As NBs and INPs continue to divide, they lose their capacity to generate early born progeny, a phenomenon termed progressive restriction in competence (Doe 2008; Farnsworth and Doe 2017). It is possible that as NBs and INPs age, they also lose proliferative potential. Perhaps *para* acts as an aide to help older cells divide, whereas *para* function may be unnecessary in younger, more robust stages of NB/INP development (Supplemental Fig. S8A-B).

Aside from development, VGSCs are widely expressed in various cancers including breast, cervical, prostate, colon, gastric, leukemia, lung and gliomas (Patel and Brackenbury 2015). Within this wide range of cancers, VGSC contributes to proliferation, metastasis, cell migration and invasion (Patel and Brackenbury 2015; Brackenbury et al. 2008). We found that *para* not only influences neural development, but is also important for tumor progression in multiple *Drosophila* brain tumor models. Knockdown of *para* in tumor tissue or introduction of a temperature sensitive hypomorphic mutant *para*^TS1^ both led to reduction in brain tumor size. As we identified roles for VGSCs in cancer progression, VGSCs represents a promising target for therapeutic intervention. VGSCs are druggable targets, and a number of FDA approved medications targeting VGSCs for pain sensation, antipsychotics and seizures currently exist in the market (Patel and Brackenbury 2015).

Our studies have identified a role for voltage-gated sodium channel, *para*, in neural development and cancer. Going forward it will be interesting to discover how the activity of other ion channels, exchangers and transporters converge to convert electrical signals into biochemical and genetic pathways to influence cellular behavior. Our findings indicate that the actions of *para* are cell autonomous, but it will be interesting to elucidate signaling between members of a cellular niche for the neuroblast lineages. Many progenitors and stem cells are connected through gap junctions and electrical signaling likely facilitates communication and coordination, essential for tissue patterning and cellular morphogenesis during development and regeneration. Genetically tractable model systems like *Drosophila* provide a platform to build this knowledge and further our understanding of the contribution of ion channels to proliferation and differentiation, in various cellular contexts. These studies will offer significant insight towards designing biomedical therapies for birth defects, cancer and regenerative medicine.

## Materials and methods

### *Drosophila* stocks and culture

*insc-Gal4* (Gal41407 inserted in inscuteable promoter) UAS-dpn for ectopic/over-expression of Dpn (Wallace et al. 2000; Zhu et al. 2012) *insc-Gal4* was used to drive the expression of transgenes in NBs. *ase-Gal80* was used to restrict the expression of transgenes to type II NB lineages (Zhu et al. 2012). UAS-*para* RNAi (BL31676 y1 v1; PTRiP.JF01469attP2) or y1 sc* v1 sev21; PTRiP.HMS00868attP2 and control stock y1 v1; PUAS-GFP.VALIUM10attP2 (BL35786) or y1 v1; PUAS-LUC.VALIUM10attP2 (BL35788) were crossed to w* UAS-mCD8-GFP; *insc-Gal4*,UAS-dcr2/CyO, tubG80; UAS-Dpn. Or w*,UAS-mCD8-GFP; r9d11-Gal4. The *para*ts (Ganetzky 1984) temperature sensitive allele was examined in type II neuroblast tumors using females *para*^TS1^; *insc-Gal4*, UAS-GFPNLS/CyO, weep, and as a control w*; insc-GAL-4,UAS-GFPNLS/CyO, weep, crossed to males UAS-dpn/TM6B, tb or *para*^TS1^;UAS-dpn/TM6B,Tb, or activated Notch designated NIC (BL5830) w-, UAS-NB2A2or brat RNAi knockdown tumor: y1 sc* v1 sev21; PTRiP.HMS01121attP2. MARCM analysis was performed with the following stocks: w* HS-FLP FRT19A; *insc-Gal4*, UAS-mCD8-GFP or w-HS-FLP FRT19A; *insc-Gal4*, UAS-GFPNLS or w-HS-FLP FRT19A; *insc-Gal4*, UAS-td-tomato crossed to control w* FRT19A or mutants w* *para*^-/-^ FRT19A/ FM7, Kr-Gal4, UAS-GFP. For rescue experiments we used w-*para*^-/-^ FRT19A/ FM7, Kr-Gal4, UAS-GFP; UAS-*para* (UAS-*para*, also known as UAS-DmNaV, was a generous gift from Marc Tanoyue’s lab), and w* *para*^-/-^ FRT19A/ FM7, Kr-GAL4, UAS-GFP; UAS-*para*D388N/CyO, Kr-Gal4, UAS-GFP. What we refer to as *para*V1401E is reported in flybase as: y1 w* *para*B/ FM7, Kr-GAL4, UAS-GFP (BL57109). FUCCI analysis was performed in stocks made from w1118; PUAS-GFP.E2f1.1-23032PUAS-mRFP1.NLS.CycB.126619/CyO, Pen1wgen11; MKRS/TM6B, Tb1 (BL55121) here shortened to FUCCI: virgin females w1118 FRT19A; FUCCI/CyO,weep or w1118*para*^-/-^FRT19A; FUCCI/CyO, Weep crossed to male w* HS-FLP FRT19A; *insc-Gal4*. Histone live imaging was performed using w1118 FRT19A; PHis2AvT:Avic-S65T62A/CyO, Kr-GAL4, UAS-GFP or w* *para*^-/-^ FRT19A/ FM7, Kr-GAL4, UAS-GFP; PHis2AvT:Avic-S65T62A/CyO, Kr-GAL4, UAS-GFP (His-GFP from BL5941) crossed to male w* HS-FLP FRT19A; *insc-Gal4*, UAS-td-tomato.

### Generation of *para* null allele

PBacWHf04029 (Exelixis:f04029 at Harvard Medical school) and PXP*para*d04188 (Exelixis:d04188 from Harvard Medical School) stocks were used to generate a deletion of the Para gene region. FLP recombinase and FRT-bearing insertion was used to generate an isogenic deletion with molecularly defined endpoints described in (Parks et al. 2004). The Exelixis deletion series is based on transposon insertions containing FRT sites of 199 bp 5 (in XP and WH transposons) of the white+ transgene. Using a heat shock FLP recombinase, trans-recombination between FRT elements resulted in a genomic deletion with a residual, hybrid element WH:XP, tagging the deletion site. Progeny were screened for the presence of residual element by PCR using WH or XP element specific primers. Absence of deleted region in null animals was examined using primers from within the gene region. Primers used for Para deletion of hybrid PCR of WH5’ plus XP5’ minus elements include forward primer: GACGCATGATTATCTTTTACGTGAC and reverse primer: AATGATTCGCAGTGGAAGGCT to generate a 1.8kb PCR band. To assess loss of Para gene region, a reverse primer within an intron: CTGCTGTATTC-GAGTCATTGG and a forward primer within a coding region: TTCGGATGGGCTTTCCTGTC generate a 500bp band.

### MARCM clones

Mosaic analysis with a repressible cell marker (MARCM) neuroblast clones were generated similar to previously described (Lee et al. 1999). *Drosophila* melanogaster were grown on standard media at 25°C. To induce mitotic recombination, staged larvae, which hatched within a 2-hour interval, were collected into vials containing about 10 mL of regular fly food at the density of approximately 80 larvae per vial, heat shocked in a 37°C water bath for 1 hour and then returned to 25°C. Dissections occurred at 48, 72 and 96 hours after larval hatching. In addition to HS-FLP recombinase, we also used an enhancer-trap recombinase, FINGR-FLP (ET-FLP 200c) (Bohm et al. 2010), which we find is sufficient to generate type I and type II MARCM clones (Figures 2A-D). The heat shock step was not necessary for this enhancer driven FLP, but collection and dissection times remained consistent to the methods described above.

### cDNA Para over expression analysis

UAS-*para* and UAS-*para*bss were gifts from Mark Tanoyue’s lab(Parker et al. 2011). UAS-*para*D388N was generated by introducing the D388N point mutation into a wildtype cDNA received as a gift from Ke Dong’s lab at Michigan State(Olson et al. 2008). The *para*D388N cDNA fly stock was generated by insertion into the VK37: (2L) 22A3 PhiC31 site.

### *Drosophila* immunohistochemistry and microscopy

Larval brains were dissected, fixed and stained similarly to previously described(Zhu et al. 2012). Briefly, third instar larvae were dissected in PBS (phosphate buffered saline), fixed in 4% formaldehyde solution for 20 min on ice, washed 3 times in PBS. Blocked for one hour in blocking buffer (PBST (PBS, 0.03%TritonX-100) with 10% normal goat serum) and incubated with primary antibodies in blocking buffer overnight at 4 °C. The following day primary antibody was removed, brains were washed 3 times with PBSTX, followed by secondary antibody incubation for 2 hours at room temperature. Primary antibodies include: guinea pig anti-ase (1:1000), rabbit anti-dpn (1:500) rat anti-cd8 gfp (1:400 from invitrogen 13-0081-82), rat anti-tdtom (Kerafast EST203). For FUCCI staining: chicken anti-gfp (1:400 from Aves lab GFP-1020) chicken anti-mcherry (1:400 from Novus biologics NBP2-25158). Cleaved Caspase staining (rabbit anti-cleaved DCP-1 (1:200; Asp216, Cell Signaling Technology 9578S). Secondary antibodies conjugated to Alexa Fluor 488 (Invitrogen A-11006) Alexa Fluor 555 (A-21428) or 633(A-21105) (Invitrogen) were used at 1:400 and DAPI staining at 1:1000. Before imaging brains were mounted onto slides, oriented, vacuum grease was placed on 4 corners and covered by a coverslip, vectashield was added and coverslip sealed with nail polish. Figure 1 brain tumor images were acquired with Leica SP5 confocal microscope with 1µm stacks. The brain lobe size was measured by Imaris 5.5 software after three-dimensional reconstruction of the z-stack of confocal images of gfp staining to specifically measure the tumor cells. For Neuroblast (NB) quantification, type I NBs were identified by Dpn+; Ase+ labeling on the surface of brain lobe excluding optical lobe region, and type II NBs were identified by Dpn+; Ase labeling on the dorsal region of brain lobe. Cellular subtypes were identified and counted as those within cellular lineages or MARCM clones marked by mCD8-GFP or nls-GFP and further classified by staining where intermediate neural progenitors are marked by Dpn+Ase+, GMCs by Ase+ and neurons by position and absence of antibody staining. MARCM clones and type II lineages for overexpression experiments were performed on were imaged on a Leica SP5 or Leica SP8 confocal microscope with 0.5µm stacks.

### Edu labeling and staining

EdU (5-ethynyl-2-deoxyuridine) labeling protocol was derived from (Daul et al. 2010). Briefly, 72hour ALH staged larvae were placed on kankel-white medium containing 0.2mM EdU and bromophenol blue for 4 hours. After 4 hours half of the larvae were dissected for a time = 0hours of EdU timepoint in schneiders medium on ice. 500uL of fix solution (4%(v/v) formaldehyde, 0.1M PIPES pH6.9, 0.3%v/v Triton x-100, 20mM EGTA pH 8.0, 1mM MgSO4) for 23 minutes at room temperature. Brains were washed 3 times in PBSTX for 20minutes at room temperature. Samples were then blocked in blocking solution (PBSBTX (1x PBSTX + 1%w/v BSA) 5uL of normal goat serum, 1M glycine made fresh and kept on ice). Click-iT™ Plus EdU Alexa Fluor™ 647 Imaging Kit was used as described in manufacturor’s protocol (Thermo Scientific C10640). 150uL of Click-iT reaction mix was prepared in the dark and let to sit no more than 5 minutes before being added to samples for 45 minutes rocking at room temperature. Brains were then washed twice in PBSTX at RT then in PBSBTX before adding primary antibody solution diluted in blocking buffer rat anti cd8 (1:400), rabbit anti Dpn (1:500) and incubating overnight at 4 degrees C. Secondary labeling continued the next day as described above.

### Xenopus Oocyte recording and RNA preparation

cRNA was isolated similar to described in (Olson et al. 2008). Briefly, 1ug of PGH19 plasmid DNA containing the DmNaV (a gift from Ke Dong’s Lab (Olson et al. 2008)) insert was linearized using the NotI restriction enzyme. The linearized DNA was used for in vitro synthesis of the DmNaV cRNA using in vitro transcription with T7 polymerase (mMESSAGE mMACHINE kit, Ambion AM1344). For robust expression of DmNaV sodium channels, DmNaVcRNA (0.25–10 ng) was co-injected into oocytes with the D. melanogaster tipE cRNA (1:1 molar ratio), which enhances the expression of insect sodium channels in oocytes (Feng et al. 1995; Warmke et al. 1997). Oocytes were harvested from the abdominal cavities of female Xenopus laevis frogs in accordance with protocols approved by the Institutional Animal Care and Use Committee at UCSF. Following manual se*para*tion of lobes and defolliculation, oocytes were injected with 50 nL of solution containing 200 ng/uL of mRNA encoding wild-type or mutant *para* as well as 200 ng/uL of tipE mRNA. Oocytes were subsequently maintained by incubation with gentle agitation at 18oC in ND96 containing, in mM: 96 NaCl, 3 KCl, 1 MgCl2, 2 CaCl2, and 5 HEPES (pH 7.4/NaOH) supplemented with 100 U/mL penicillin, 100 µg/mL streptomycin, and 50 µg/mL gentamicin until the day of recording. ND96 was exchanged twice daily during incubation. Two Electrode Voltage Clamp Electrophysiology and Data Analysis 48-72 hours after injection, oocytes were transferred individually to a recording bath containing ND96 (not supplemented with antibiotic) at room temperature, and impaled with two borosilicate glass electrodes (tip resistance 0.2-1.0 MO) filled with 3 M KCl. Recordings were made using a GeneClamp 500B amplifier and Digidata 1320A digitizer (Axon Laboratories) driven by pClamp10 software. After successful establishment of membrane voltage clamp, individual sweeps were collected at a 20 kHz sampling rate and subjected to 6x P/N subtraction online, and were then low-pass filtered off-line using an 8-pole Bessel filter at 2 kHz. All recordings were made from a holding potential of −80 mV. Non-normalized anti-peak values were collected from each sweep and averaged to generate I/V relationships, and are presented as mean +/− S.E.M., with n values representing total number of individual cells. All constructs were tested on at least 4 separate days of recording.

## ACKNOWLEDGEMENTS

We thank Ruijun Zhu, Ke Li, Caitlin O’Brien, Han-Husan Liu, Jacob Jaszczak, David Crottes, Maja Petkovic, Mu He, Chin Fen Teo, Cindy Li and other Jan lab members for helpful discussions and experimental consultations. We thank Mark Tanoyue (UC Berkeley) for generous gifts of fly stocks and Ke Dong (Michigan State University) for generous gifts of *para* cDNA and helpful guidance for culturing conditions. Stocks obtained from the Bloomington *Drosophila* Stock Center (NIH P40OD018537) were used in this study. Funding was provided by a Lefkofsky foundation Damon Runyon Cancer Research Foundation Fellowship (B.J.P) and NIH grant 1R35NS97227 to Y.N.J.. Y.N.J. and L.Y.J. are investigators at the Howard Hughes Medical Institute

---

**Supplemental Fig. S1.**
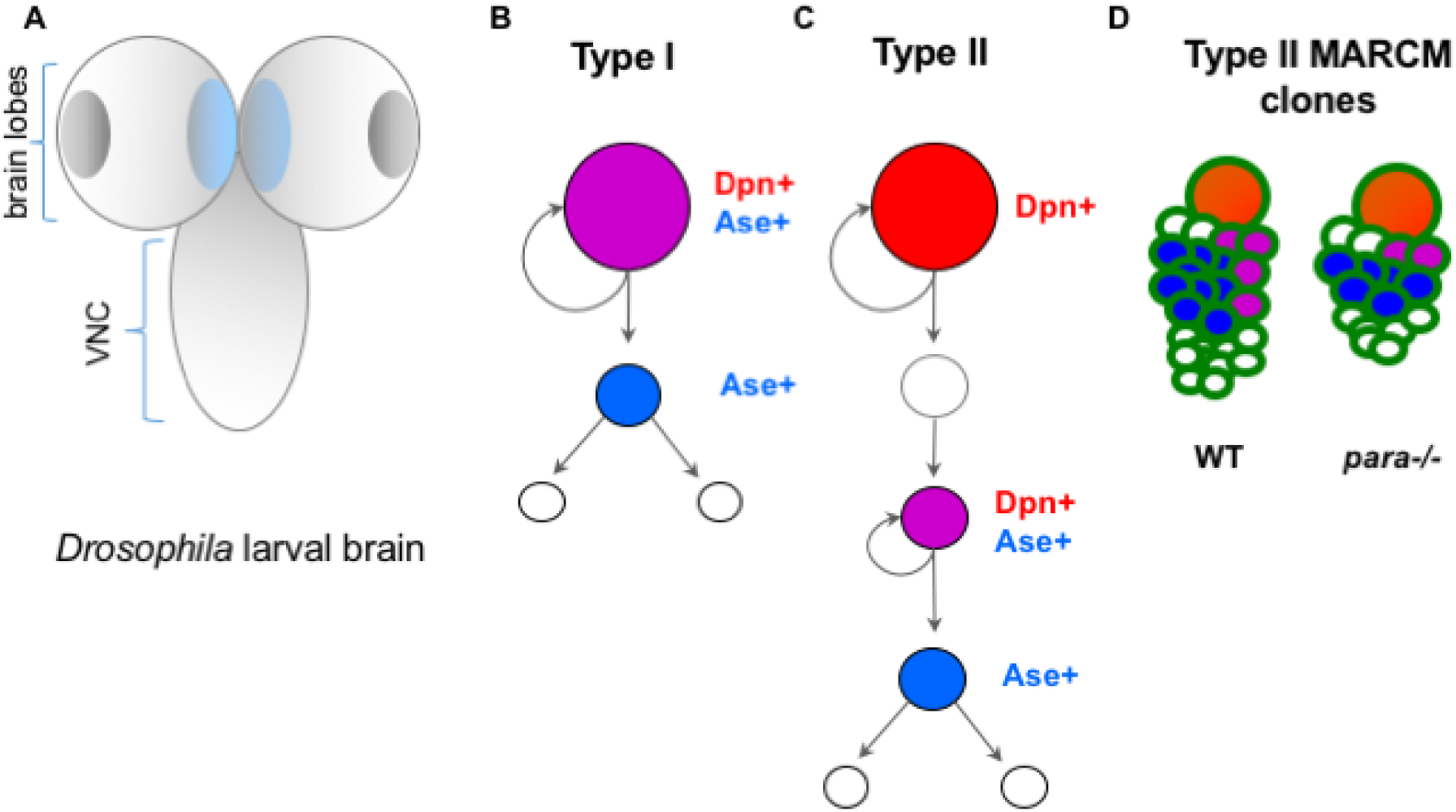
Cartoon schematic of *Drosophila* larval brain development. **(A)** Cartoon of *Drosophila* larval brain consisting of 2 brain lobes and a ventral nerve cord. Darker grey areas indicate the optic lobe while lighter grey regions are where type I neuroblasts (NBs) reside. Light blue area within the central brain lobes indicates regions of type II neuroblasts. **(B)** Type I pattern of division. Type I NBs (In magenta, Deadpan^+^ (Dpn) and Asense^+^ (Ase)) undergo asymmetric divisions to self-renew and generate a more differentiated Ase^+^ ganglion mother cell (GMC) in blue. The GMC will go on to symmetrically divide into two neurons or glia. **(C)** Type II NBs (in red) are Dpn+Ase-. They asymmetrically divide to self-renew and generate an immature intermediate neural progenitor (INP). This INP will mature (in magenta) and become Dpn^+^Ase^+^ and go on to asymmetrically divide to self-renew and generate a more differentiated GMC (blue), which will symmetrically divide to generate two neurons or glia. **(D)** Cartoon images of type II lineage MARCM (Mosaic Analysis with a Repressible Cell Marker) wildtype (WT) and *para*^*-/-*^ dones display NB and progeny. Earlier bom progeny with more proliferative potential are closer to NB while later born, more differentiated progeny are further away from the NB. *para*^*-/-*^ MARCM clones possess all type II progeny cell types, but in fewer numbers compared to WT.

**Supplemental Fig. S2.**
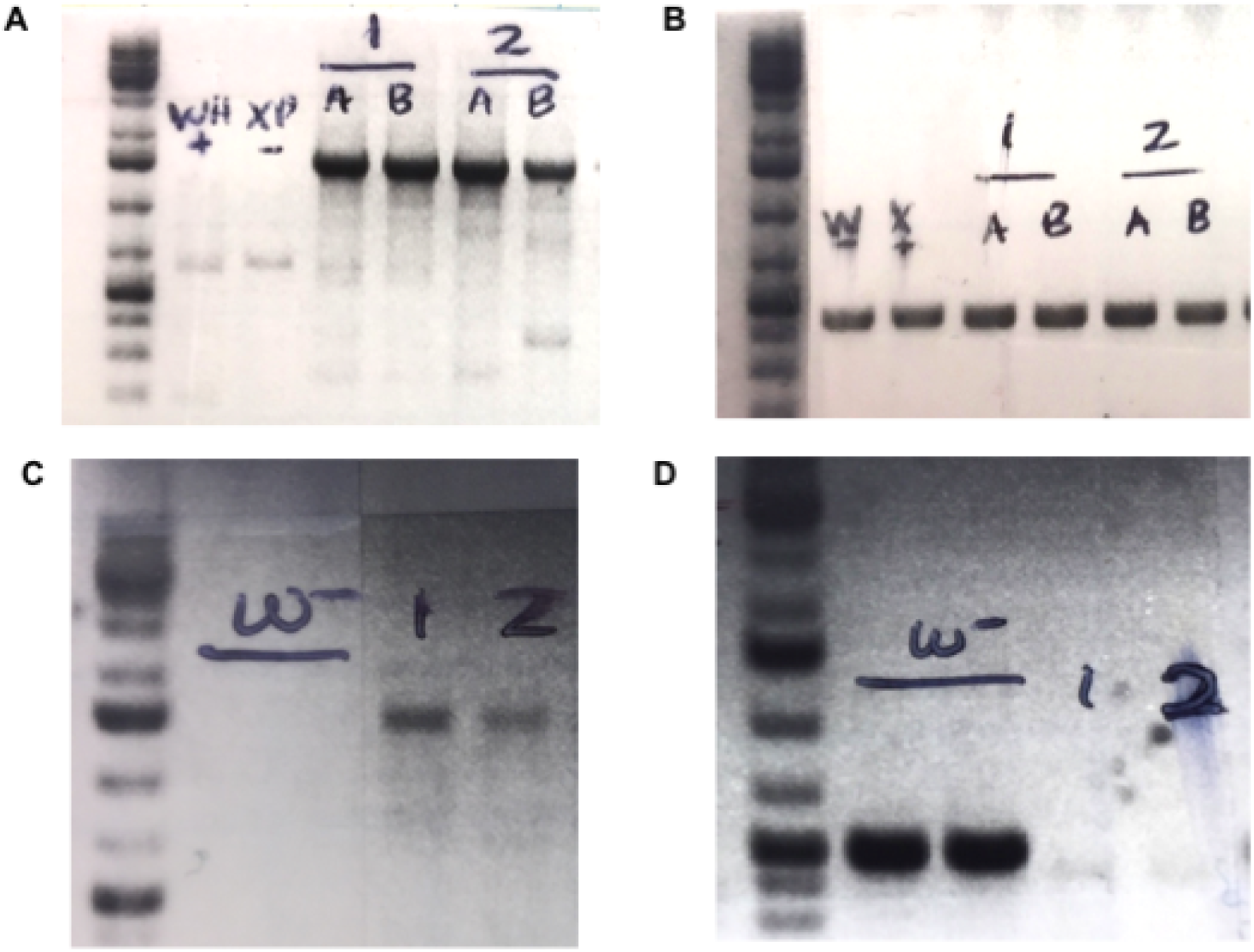
Generation of *para*^*-/-*^ allele. DNA gel showing genotyping of wildtype and *para*^*-/-*^ alleles. Genomic DNA was isolated from a collection of approximately 10 larvae. **(A)** hybrid PCR generated by primers flanking deletion. In para-/- animals labeled 1 and 2, a 1.8kb band of DNA is amplified by PCR. Wildtype stocks denoted WH(**+**) and XP(**-**) used to make *para-/-*, there is no 1.8kb band as the aene region between the flanking primers is approximately 63kb and does show up on the gel. **(B)** Internal gene PCR of a 500bp region found within *para* gene region. This band is present in wildtype (labeled W- and X+) as well as heterozygous fly stocks labeled 1(**A,B**) and 2 **(A, B). (C)** While loss of *para* is lethal, a small fraction of *para-/-* larvae hatch and die shortly after They are identified by absence of balancer KGB and display sluggish, uncoordinated movements. Wildtype (*w*-) first instar larvae lack 1.8kb flanking PCR band, but the band is present in 1^st^ instar *para*^*-/-*^ larvae (labeled 1 and 2). **(D)** Internal 500bp DNA band is present in wildtype larvae (w-), but lost in null mutants identified by phenotype (labeled 1 and 2).

**Supplemental Fig. S3.**
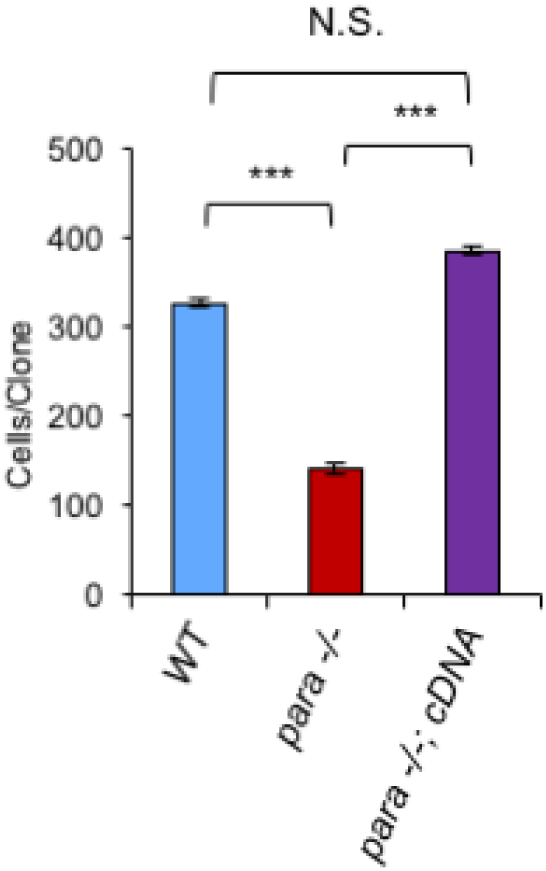
*para* cDNA rescues cell number in type 1 and type II neuroblast lineages. Similar to 72h ALH, at 96h ALH *para*^*-/-*^ reduction in clones per MARCM clone compared to WT is rescued by cell autonomous expression of a Para cDNA. One-way ANOVA with Bonferonni multiple comparison test, *** p<0.0001 WT N=14 para^*-/-*^ N=19 *para*^*-/-*^; UAS-*para* cDNA N=11 para-/- MARCM clone cell number deficit is rescued by cell autonomous expression of a *para* cDNA, time = 96h ALH. One-way ANOVA with Bonferonni multiple comparison test, ***p<0.0001 (96h ALH: WT N=31 *para*^*-/-*^ N=20 *para*^*-/-*^; UAS-Para cDNA N=6).

**Supplemental Fig. S4.**
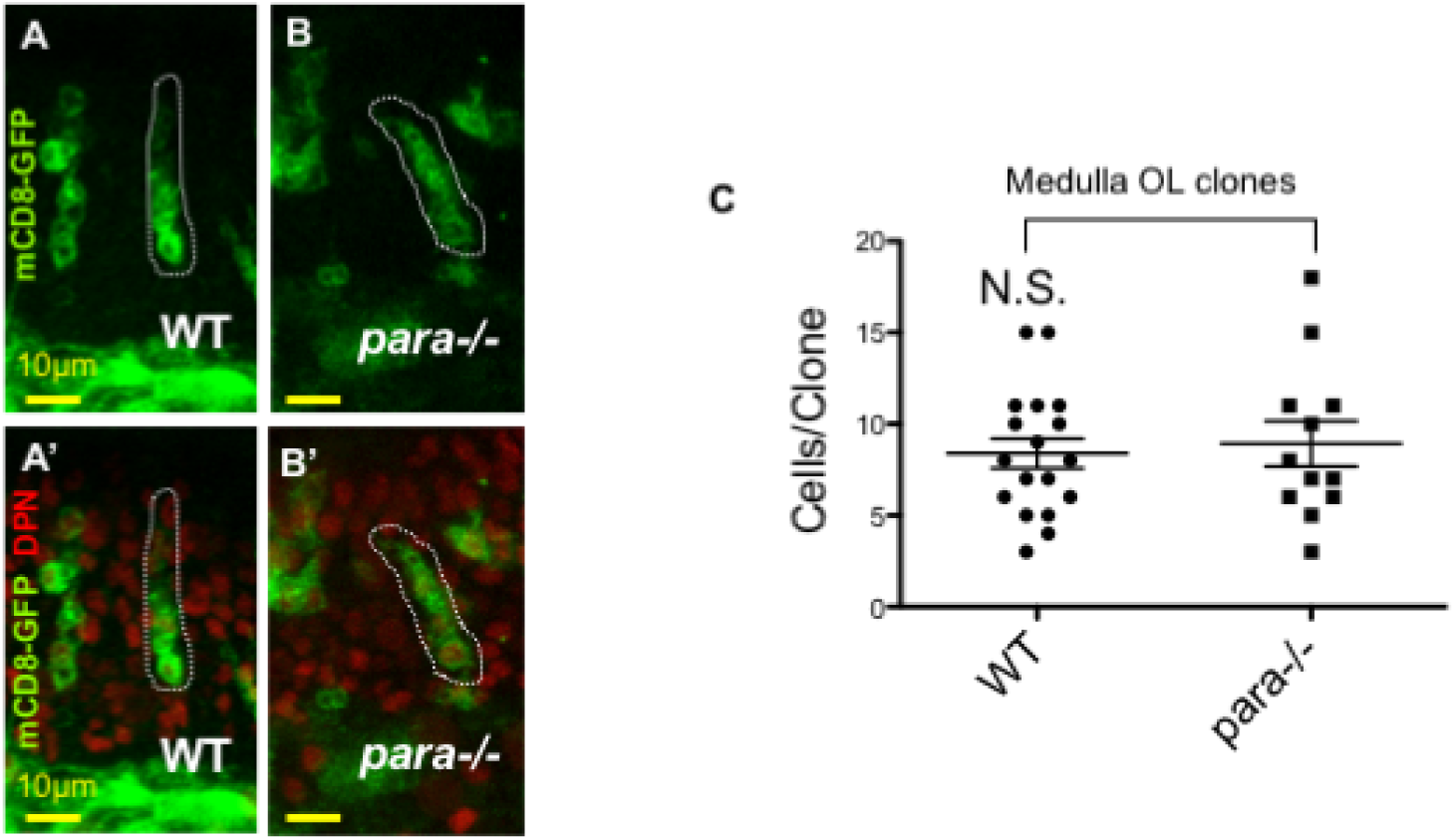
Within the optic lobe, *para*^*-/-*^ medulla neuroblast clones display similar numbers of Mils compared to WT clones. **(A-B**^**’**^) Representative images of optic lobe MARCM clones labeling medulla neuroblast lineages (driven by a heat shock (HS) FLP recombinase at 72h ALH with *insc-Gal4*, outlined in white dashed line). Scale bar =10µm. In contrast to type I and II central brain neuroblast lineages, medulla optic lobe clone numbers appear unaffected by genetic removal of *para* (**(C)** unpaired, two-tailed t-test p>0.05 WT N=18, *para*^*-/-*^ N=12).

**Supplemental Fig. S5.**
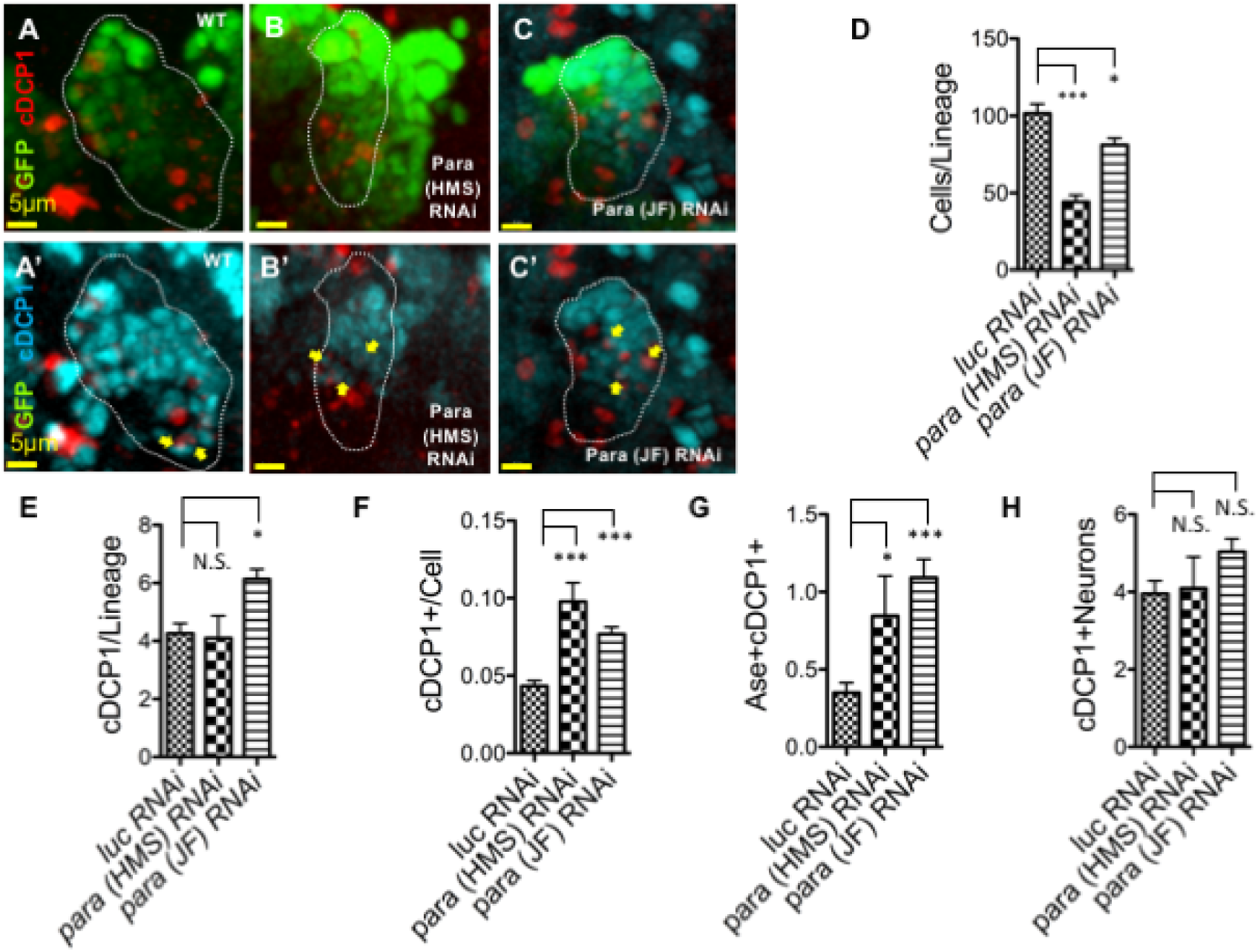
RNAi Knockdown of Para reduced cell numbers and increased cleaved caspase staining in Ase+ cells within the type II lineage. **(A-C)** Representative images nuclear localized GFP within *luciferease (lue)* RNAi control or para RNAi stocks was driven by *inscuteable Gal-4; ase-Gal80* to restrict expression to the type II lineage show reduced cell numbers in both para *RNAi* knockdown stocks, Scale bar =5μm (**(D)**, *luc RNAi* N=16, HMS N=12, JF=13, p<0.05, ANOVA with Dunnett’s multiple comparison test). **(A-C’)** Representative images showing co-labeling of cDCP1 and Ase+ cells indicated by yellow arrows (GMCs or INPs). (E-F) The weaker JF *para RNAi* showed more cDCP1 staining than control **(E,** *luc RNAi* N=16, HMS N=12, JF=13, p<0.05, ANOVA with Dunnett’s multiple comparison test), while both HMS and JF *para RNAi’s* show significant increase of cDCP1/cell when lineage size is taken into account ***((F)***,*luc RNAi* N=15, HMS N=8, JF=12, p<0.05, ANOVA with Dunnett’s multiple comparison test). **(G-H)** Both *para* RNAi’s lead to increased numbers of Ase+ cells colabeled with CDCP1+ **(G, )** *luc RNAi* N=15, HMS N=8, JF=13, p<0.05, ANOVA with Dunnett’s multiple comparison test), but there is no significant difference in neuronal cDCP1 labeling at this time point **(H, )**, *luc RNAi* N=15, HMS N=8, JF=13, p>0.05, ANOVA with Dunnett’s multiple comparison test).

**Supplemental Fig. S6.**
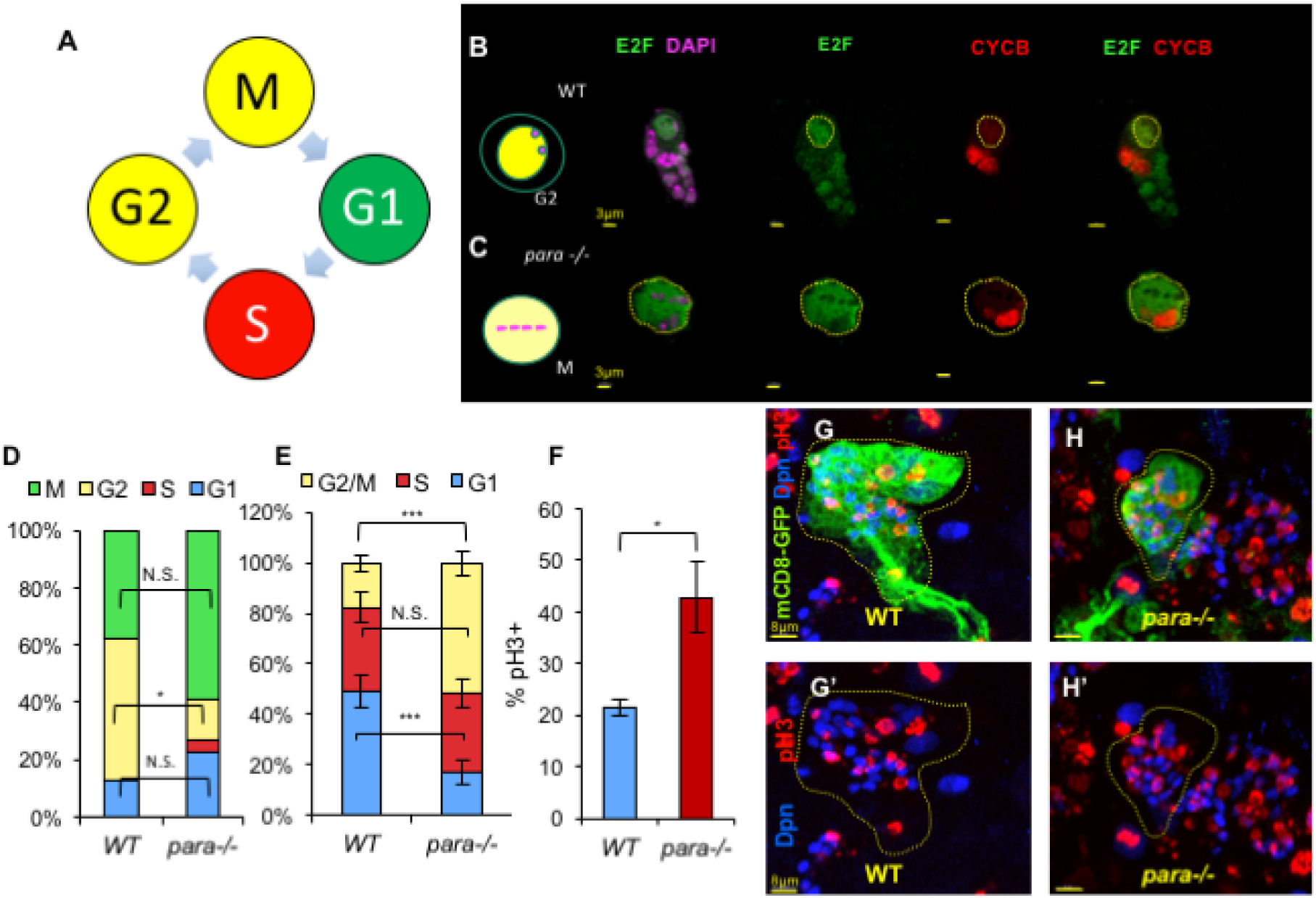
Genetically encoded cell cycle indicator FUCCI in neuroblast lineages revealed tyoe II progeny enrichment in G2/M phase. **(A)** Fly-FUCCI encodes a genetic cell cycle sensor with a nuclear localized cyclin B (CycB) sequence tagged with red fluorescent protein (RFP) and cyclin E2F sequence tagged with nuclear GFP. E2F is on during all cell cycle stages except S phase, while cydin B is on during all phases except G1.Using the combination of fluorophores one can deduce the state of the cell cycle. As M and G2 are both E2F+CycB+ they can be differentiated using DAPI to highlight chromosome condensation which occurs during mitosis. **(B-C)** Sample images showing E2F-GFP, CycB-RFP and DAPI staining. Type II MARCM clone samples WT (top) *para*^-/-^ (Bottom) Cartoon of cell in mitosis for *para*^-/-^ shows yellow cell representing E2F-GFP+, CycB-RFP+ with absence of nuclear membrane and condensed chromosomes lined up in meta phase. WT cartoon is also E2F-GFP+, CycB-RFP+, but still has intact nuclear membrane prior to M-phase and condensed chromosomes indicating the cell is in G2 prior to mitotic entry and nuclear membrane breakdown. Scale bar Is 3μm. **(D)** FUCCI analysis showed a reduced fraction of type II para, NBs (N=22) in G2 compared to wildtype NBs (N=16) (96h ALH, unpaired two-tailed t-test, P=0.0179) **(E)** A significant number of mitotically active type II *para*^-/-^ progeny (INPs and GMCs), were enriched in G2/M phase at the expense of Gl compared to WT. (96h ALH, Two-tailed T-test, p<0.0001. WT N=22 *para*^-/-^ N=24). **(F)** Due to their small size it was challenging to differentiate between G2 and M-phase using DAPI in NB progeny, thus pH3 staining was employed to examine percent of INPs in mitosis. We found para-/- INPs have an increased number of pHB-+ cells compared to WT controls. (96h ALH, WT N:8 *para*^-/-^ N=5 Two-tailed T-test, p<0.0001). **(G-H’)** Representative images of maximum projection stack of type II MARCM clones (outlined by green, membrane labeled mCD9>6FP) showed *para-/-* Dpn+ INPs (labeled in blue) have a higher proportion of pH3+ (labeled in red) than WT, Scale bar = 8μm.

**Supplemental Fig. S7.**
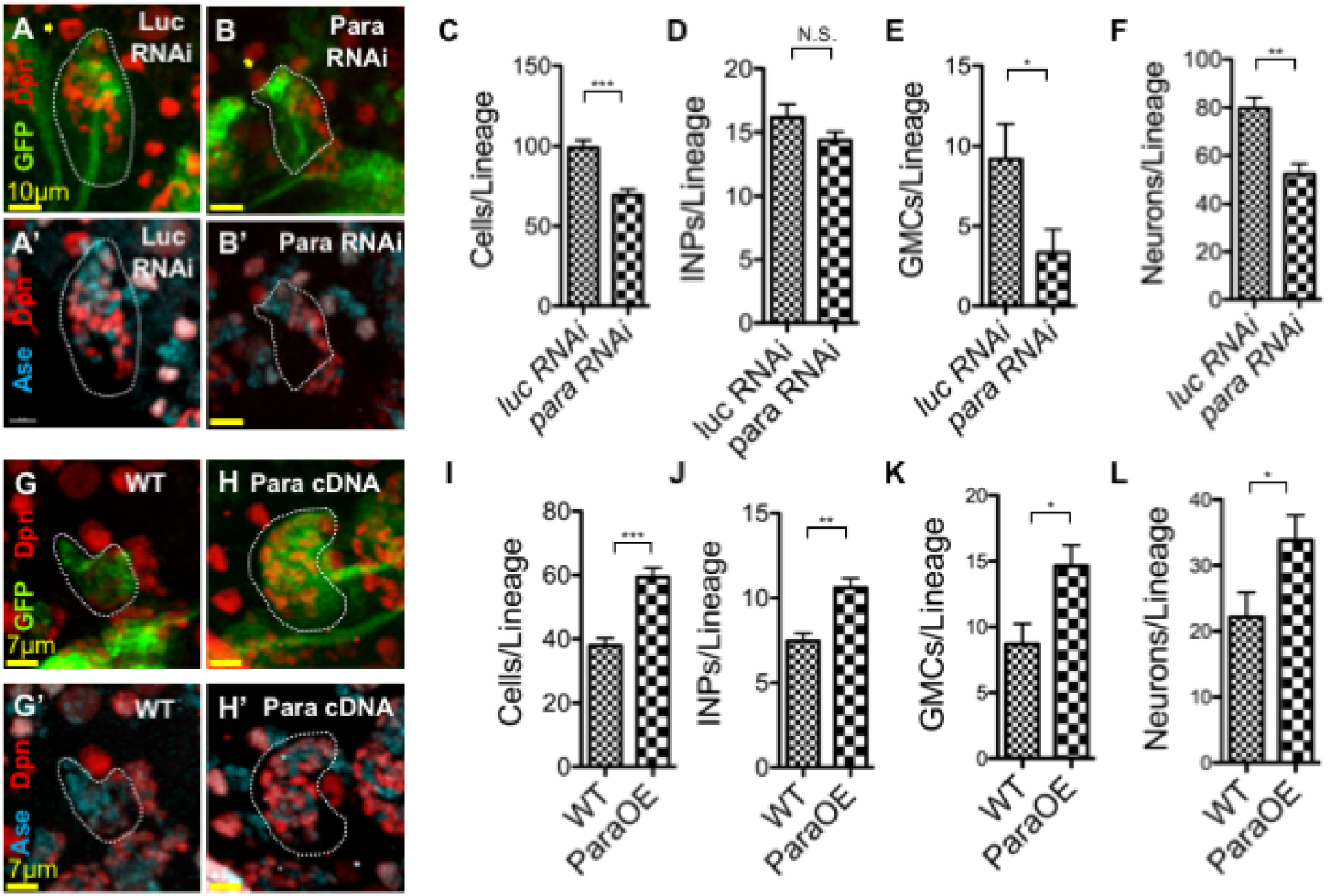
Altered exprassion level of Para driven wlttiln INPs, but not NBs, influenced cell numbers in Type II lineages. **(A-B’,C)** Representative images of RNAi knockdown of *para* driven by *r9d11-Gal4*, scale bar: 10μm **(A-A’)** only within INPs was sufficient to reduce numbers of cells per type II neuroblast lineage compared to *luciferase (luc)* RNAi control **(B-B’)** (96h ALH, unpaired two-tailed t-test, P< 0.0001, *luc RNAi* N=14, *para (HMS) RNAi* N=14) **(D)** INP numbers were not changed with *para RNAi* knockdown, however **(E)** GMC and **(F)** neural numbers were reduced in para RNAi (Two-tailed T-test, INP p=N.S., *luc RNAi* N=14, *para RNAi* N=13, GMC p<0.05, *luc RNAi* N=7, *para (HMS) RNAi* N=14, Neurons p<0:01,*luc RNAi* N=6, *para (HMS) RNAi* N=14). **(G-I)** Representative images, scale bar = 7μm (G-H’) of overexpression of *para* (ParaOE) wildtype cDNA driven within the INPs *(r9d11-Gal4)* of the type II lineage is sufficient to increase total cells per type ill lineage as well as all cellular subtypes **(J-L).** (72h ALH, unpaired two-tailed t-test, p< 0.0001, total cells: WT N=13, ParaOE N=11, INPs, p<0.01: WT N=13, ParaOE N=11, GMCs, p<0.05: WT N=8, ParaOE N=9, neurons, p<0.05: WT N=8, ParaOE N=9).

**Supplemental Fig. S8.**
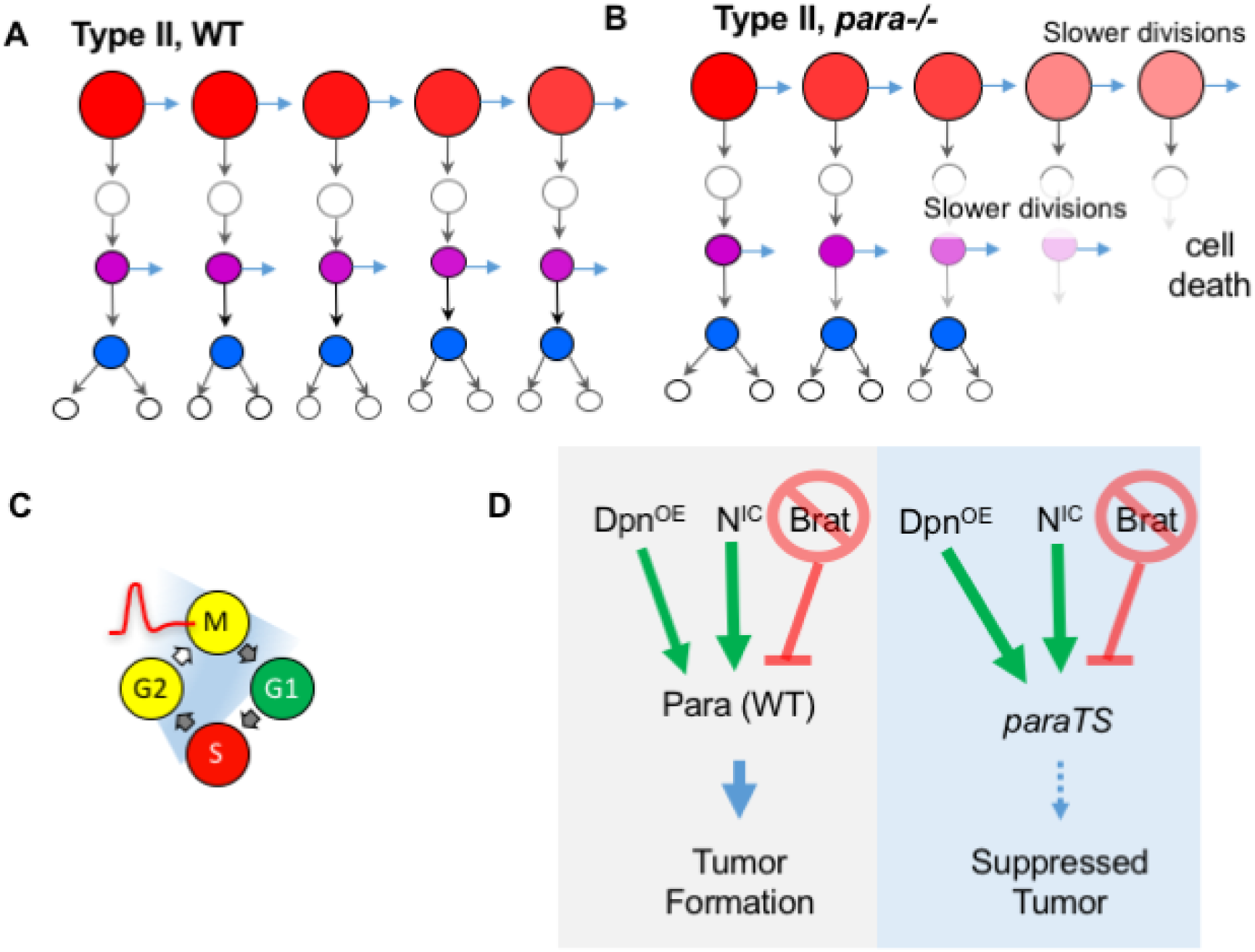
Model for Para’s role in progenitor proliferation. **(A)** Neuroblasts asymmetrically divide to self-renew and generate an intermediate neural progenitor (INP) which itself will self-renew and generate a ganglion mother cell (GMC). GMC will symmetrically divide to generate two neuron or glia. **(B)** *para*^-/-^ type II NBs and INPs may lose proliferative potential as they progressively self-renew and age. This may result in slower proliferation, fewer progeny generated and premature cell death in INPs. **(C)** Para channel activity may provide depolarization to activate downstream signaling cascades to promote G2-M transition in the cell cycle. Loss of this activity may diminish cel cycle progression. **(D)** Overexpression of stem cell promoting genes Dpn or Notch, or knockdown of cell fate determinant Brat are sufficient to generate tumor formation in neuroblast lineages. Para likely acts downstream of these pathways as reduction of Para (with hypomorph *ParaTS)* in these tumor backgrounds is sufficient for tumor suppression.

**Supplemental Fig. S9.**
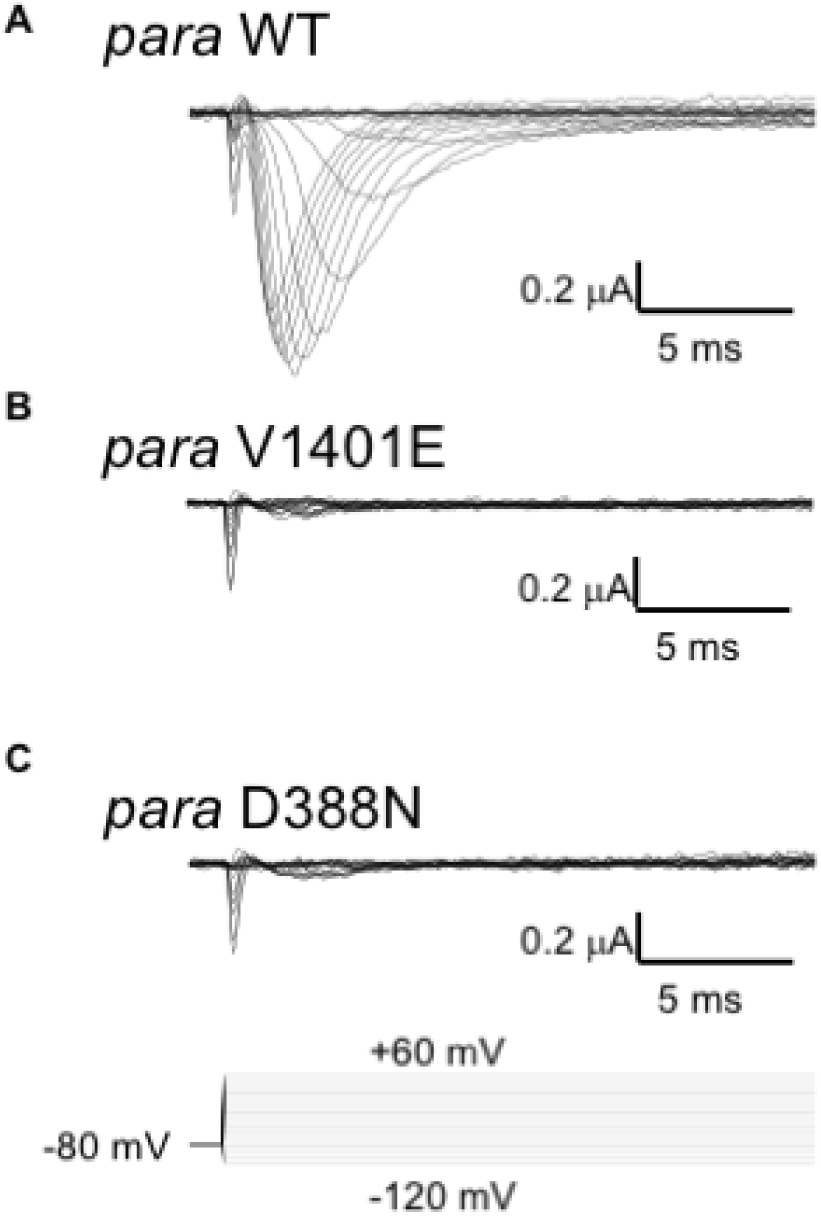
Sample Traces of Para currents in response to various changes in voltage. *Xenopus* oocytes recorded using two-electrode voltage clamp.**(A)** Wild-type *para* demonstrated a mean peak I-V at 0 mV of −0.87 +/− 0.11 mA. **(B)** Expression of the channel carrying the mutation V1401E caused a significant reduction in current amplitude, with a mean peak of −0.11 +/− 0.01 mA. **(C)** *para*^*D388N*^ demonstrated currents barely distinguishable from background with a mean peak I-V of −0.07 +/− 0.01 mA.

